# Eudicot primary cell wall glucomannan is related in synthesis, structure and function to xyloglucan

**DOI:** 10.1101/2022.05.11.491508

**Authors:** Li Yu, Yoshihisa Yoshimi, Rosalie Cresswell, Raymond Wightman, Jan J. Lyczakowski, Louis F.L. Wilson, Konan Ishida, Katherine Stott, Xiaolan Yu, Stephan Charalambous, Joel Wurman-Rodrich, Ray Dupree, Oliver M. Terrett, Steven P. Brown, Henry Temple, Kristian B.R.M. Krogh, Paul Dupree

## Abstract

The functional differences between plant cell wall hemicelluloses such as glucomannan, xylan and xyloglucan (XyG) remain unclear. These polysaccharides influence assembly and properties of the wall, perhaps by interacting with cellulose to affect the deposition and bundling of the fibrils. As the most abundant hemicellulose, XyG is considered important in eudicot primary cell walls (PCWs), but plants devoid of XyG show relatively mild phenotypes. We report here that a patterned β-galactoglucomannan (β-GGM) is widespread in PCW of eudicots and shows remarkable similarities to XyG. The sugar linkages forming the backbone and side chains of β-GGM are analogous to those that make up XyG, and moreover, these linkages are formed by glycosyltransferases from the same CAZy families. Solid-state NMR indicated that β-GGM shows low mobility in the cell wall, consistent with interaction with cellulose. Although Arabidopsis β-GGM synthesis mutants show no obvious growth defects, genetic crosses between β-GGM and XyG mutants produce exacerbated phenotypes compared to XyG mutants. These findings demonstrate a related role of these two similar but distinct classes of hemicelluloses in PCWs. This work will provide new avenues to study the roles of both β-GGM and XyG in PCWs.

**One sentence summary:** Patterned β-GGM resembles xyloglucan in structure, biosynthesis and function.

## Introduction

Although the primary cell wall (PCW) is strong enough to protect the plant cell from osmotic lysis and to maintain cell and tissue shape, it can also allow the cell to expand irreversibly during growth. How the cell wall accommodates both these contrasting and fundamental properties is poorly understood. The PCW is a composite of relatively rigid cellulose microfibrils embedded in a highly hydrated matrix of non-cellulosic polysaccharides. The hemicellulose polysaccharides xyloglucan (XyG), xylan and glucomannan are able to bind tightly to cellulose (Cavalier et al., 2008; Cosgrove, 2014; Simmons et al., 2016; Terrett et al., 2019). For many years a cellulose-XyG network was proposed to be the principal load-bearing structure of the PCW in dicots (Cosgrove, 2018). However, experimental data is now more consistent with a view where cellulose fibril interactions largely determine wall extensibility (Zhang et al., 2021). Hemicelluloses such as XyG may influence cell wall extensibility through binding at potential localised sites of cellulose fibril interaction (hot spots) (Park and Cosgrove, 2015). How the different hemicelluloses contribute to plant cell wall assembly remains an important challenge in cell wall biology.

XyG is the best studied PCW hemicellulose with a repeating patterned structure. In most dicots, this unit is normally comprised of four β-1,4-linked glucosyl residues (Glc), with the first three backbone residues in each unit substituted with α-1,6-xylosyl (Xyl) branches. This unit can be conveniently described as ‘XXXG’ (Supplemental Figure S1) using the established nomenclature (Fry et al., 1993). Xyl residues at positions two or three can be further decorated with β-1,2-galactose (e.g. XXLG), galacturonic acid or a variety of other sugars (Pauly and Keegstra, 2016), some of which may be further decorated with α-1,2-fucose. The XyG side chains probably influence the solubility of the polysaccharide during synthesis and secretion, as well as in the cell wall (Whitney et al., 2006; Han et al., 2020). The importance of the repeating structure of XyG is unclear, but it may influence how XyG adheres to surfaces of cellulose, impacting PCW properties (Zhao et al., 2014; Park and Cosgrove, 2015; Benselfelt et al., 2016). Indeed, the regular pattern of substitution of xylan, an unrelated hemicellulose, is thought to influence the binding of xylan to cellulose in secondary cell walls (Simmons et al., 2016; Grantham et al., 2017). The complete loss of XyG in the *xxt1 xxt2* Arabidopsis xylosyltransferase mutant affects the production and arrangement of cellulose in PCW in hypocotyls (Xiao et al., 2016; Zhao et al., 2019). However, this XyG mutant, and also the XyG-deficient quintuple *cslc* backbone synthesis mutant, have just small perturbations in growth (Cavalier et al., 2008; Kim et al., 2020), raising questions about the importance of this hemicellulose in PCW. In contrast, the loss of MUR3-dependant β-1,2-galactosylation results in a ‘cabbage-like’ rosette and dwarfed growth (Tamura et al., 2005; Tedman-Jones et al., 2008). This reveals a specific and important role of the XyG disaccharide side chain (and its fucosylated derivative), which may maintain XyG solubility during secretion or assembly of the wall (Aryal et al., 2020; Velasquez et al., 2021).

In the glucomannan of secondary cell walls (SCWs), the backbone of β-1,4-linked mannosyl (Man) residues is randomly interspersed with β-1,4-Glc residues and sometimes bears occasional α-1,6-linked galactose (Gal) branches. The Man residues are often acetylated (we refer here to this hemicellulose as acetylated galactoglucomannan, AcGGM) (Goubet et al., 2009; Scheller and Ulvskov, 2010; Rodrı’guez-Gacio et al., 2012). Such glucomannans are particularly abundant in gymnosperm SCW, where they interact with cellulose (Terrett et al., 2019; Cresswell et al., 2021). However, in contrast to the random backbone of the AcGGM polymer, a glucomannan from Arabidopsis seed mucilage has recently been found to exhibit a repeating backbone of the disaccharide [4-Glc-β-1,4-Man-β-1,], with frequent α-1,6-Gal branches on the Man residues (Voiniciuc et al., 2015; Yu et al., 2018). A glucomannan with elements of this repeating backbone has been reported from kiwifruit and tobacco cell cultures (Sims et al., 1997; Schröder et al., 2001), but the structure of PCW glucomannan is, in general, not well characterised.

Evidence for the importance of glucomannan in the PCW has been obtained from mannan biosynthesis mutants. The α-1,6-Gal substitutions on the glucomannan of Arabidopsis mucilage are added by Mannan Alpha Galactosyl Transferase 1 (MAGT1)/MUCILAGE-RELATED10 (MUCI10) in CAZy family GT34 (Voiniciuc et al., 2015; Yu et al., 2018). Mutants in this glucomannan galactosylation show defective mucilage architecture and cellulose rays. CSLA enzymes from CAZy family GT2 synthesise the glucomannan backbone (Liepman et al., 2005; Liepman et al., 2007). The mucilage glucomannan backbone is made by CSLA2, and *csla2* mutants also show defective mucilage architecture (Yu et al., 2014). Arabidopsis mutants in CSLA9, which is largely responsible for SCW glucomannan synthesis, show no obvious changes in wall properties (Goubet et al., 2009). However, the embryo lethality of the Arabidopsis *csla7* mutant suggests an important role of glucomannan, at least in embryonic PCWs (Goubet et al., 2003). Recently glucomannan has also been implicated in etiolated hypocotyl gravitropic bending, which involves asymmetric cell expansion (Somssich et al., 2021). Despite these examples that glucomannan is important in some instances, the role for PCW glucomannan in plant growth and development, and whether that role is related to that of other hemicelluloses, remains obscure.

Here, we investigate the structure, synthesis and function of PCW glucomannan. We report that a novel type of mannan is widely present in eudicot PCWs, and we name it <u>β</u>-<u>G</u>alacto<u>G</u>luco<u>M</u>annan (β-GGM). The β-GGM has a repeating backbone structure with evenly spaced α-Gal substitutions, some of which are further substituted with β-1,2-Gal. We identify the biosynthetic machinery required to synthesise the backbone and sidechains. β-GGM has not only many structural and biosynthetic similarities with XyG, but it may also share some functions with XyG in the PCW. These results demonstrate that distinct hemicelluloses can have associated functions and that a patterned PCW hemicellulose in addition to XyG may have importance for cell expansion and plant development.

## Results

### Two glucomannan types with distinct structures, synthesised by CSLA2 and CSLA9, are widely present in Arabidopsis PCW-rich tissues

We recently found that Arabidopsis mucilage galactoglucomannan has a structure distinct from SCW acetylated glucomannan (AcGGM) (Yu et al., 2018). We therefore hypothesised that the fine structure of PCW glucomannan might also be distinct from SCW glucomannan. To investigate this, we digested alkali-extracted cell walls from etiolated Arabidopsis seedlings (which have relatively little tissue with SCW) with mannanase *Cj*Man26A, which cleaves galactoglucomannan, yielding products with an unsubstituted Man residue at the reducing end (Gilbert, 2010; Yu et al., 2018). Using polysaccharide analysis by carbohydrate electrophoresis (PACE), we observed several different mannanase products (Figure 1A and Supplemental Figure S2). To determine the biosynthetic origin of these glucomannan fragments, we also analysed cell wall material from *csla2* and *csla9* mutants. Digestion of the *csla2* mutant walls released mainly oligosaccharides with a low degree of polymerization (DP), whereas the *csla9* mutant walls yielded longer oligosaccharides, with four main oligosaccharides (named S1–S4). In contrast, mannanase digestion of the *csla2 csla9* double mutant walls released almost no detectable oligosaccharides. These results show that CSLA2 and CSLA9 are together necessary for the synthesis of most *Cj*Man26A-digestible glucomannan in seedlings, and that each CSLA enzyme synthesizes glucomannans with distinct structures.

**Figure 1.**
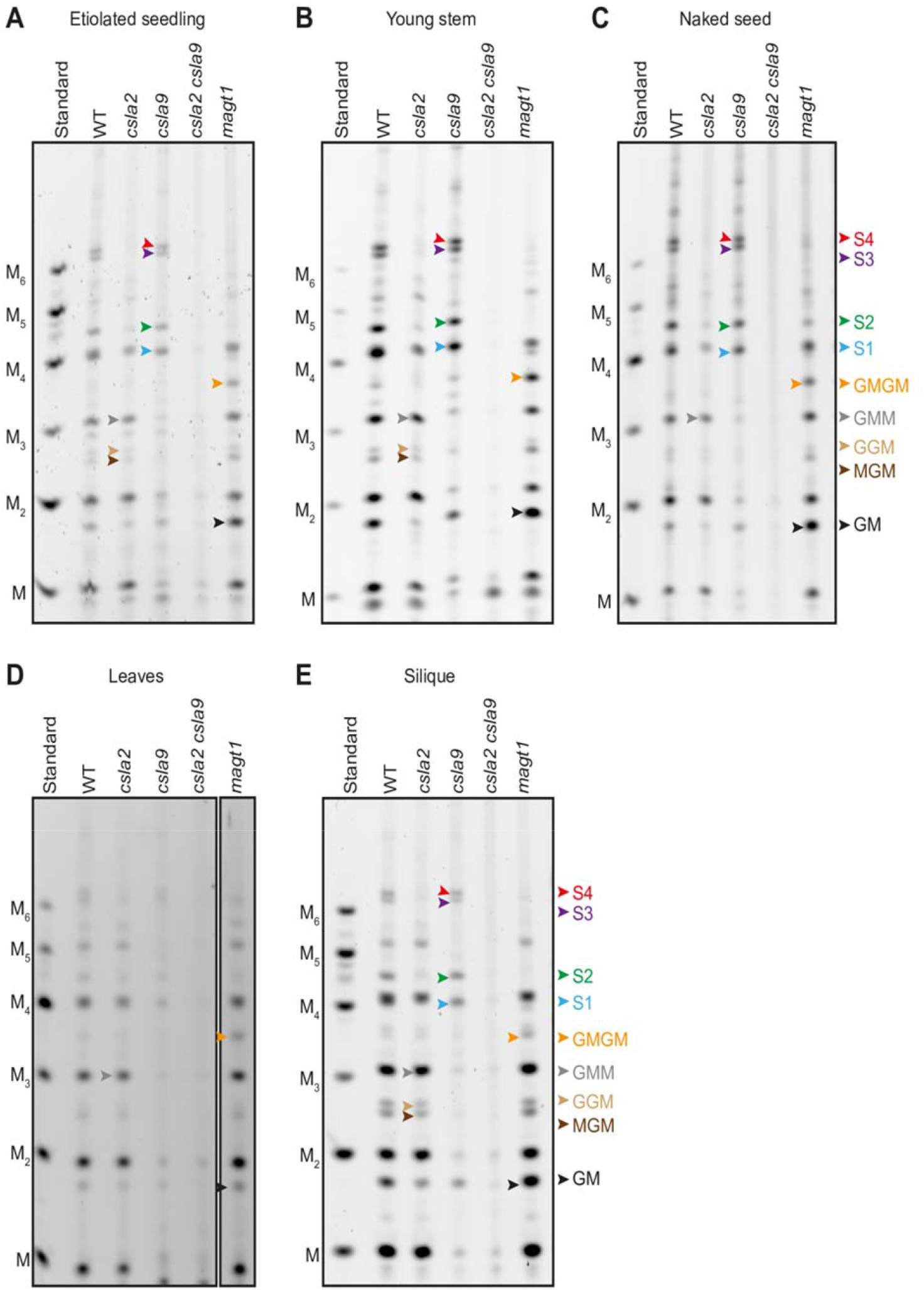
Two glucomannan types with distinct structures, synthesised by CSLA2 and CSLA9, are widely present in Arabidopsis PCW-rich tissues. Materials from five tissues, comprising etiolated seedling, young stem, seeds with mucilage removed (naked seed), leaves, and silique, were analysed by PACE. Hemicelluloses were extracted from Col-0, *csla2, csla9, csla2 csla9*, and *magt1* cell wall material using alkali before being hydrolysed with *endo*-mannanase *Cj*Man26A. The products were subsequently derivatised with a fluorophore and separated by gel electrophoresis. The *csla2* mutant yielded oligosaccharides with a low degree of polymerization, whereas the WT and *csla9* mutant walls yielded longer oligosaccharides. The four main oligosaccharides (named S1–S4) are labelled with coloured arrows in samples from *csla9*. In leaves, the amount of S1–S4 was low, and they are missing in *magt1* mutants. M, Man; G, Glc; Manno-oligosaccharide standards M to M_6_ are shown.

To investigate the mannan present in other PCW-rich tissues of Arabidopsis, alkali-extracted cell walls from young stem, seeds with mucilage removed (naked seeds), siliques and leaves were also digested with *Cj*Man26A, and the released oligosaccharides visualised by PACE. The proportion of CSLA2- and CSLA9-dependent glucomannan oligosaccharides was similar in most of the tissues, and in each case, virtually no oligosaccharides were released from the *csla2 csla9* double mutant (Figure 1, A-E). However, in leaves, the CSLA9-dependent oligosaccharides were dominant, which suggests that CSLA9-dependent glucomannan can predominate in PCW in some tissues (Figure 1D). Together, our data indicate that two distinct glucomannans, with synthesis dependent on CSLA2 or CSLA9, are widely present in Arabidopsis PCW-rich tissues.

### <u>β</u>-<u>G</u>alacto<u>G</u>luco<u>M</u>annan (β-GGM) is a patterned glucomannan with similarities to xyloglucan

To determine the structures of the distinct glucomannan polysaccharides, we characterised the oligosaccharides released from the CSLA2- and CSLA9-dependent glucomannans. We focussed first on the CSLA9-dependent oligosaccharides from *csla2* plants. From their migration in the PACE gel, we assigned the main *Cj*Man26A products as mannose, mannobiose, Glc-β-1,4-Man-β-1,4-Man (GMM, using a single letter code for each position), and Man-β-1,4-Glc-β-1,4-Man-β-1,4-Man (MGMM), consistent with a random dispersion of Glc residues in the backbone—as reported in AcGGM from gymnosperm and angiosperm SCWs (Arnling Bååth et al., 2018). To help confirm these assignments, we treated the oligosaccharides with β-glucosidase and β-mannosidase, which can only fully depolymerise the backbone in the absence of α-Gal branches. PACE analysis of the products indicated that β-glucosidase and β-mannosidase could convert the CSLA9-dependent oligosaccharides to monosaccharides and disaccharides (Figure 2A). Hence, we could deduce that CSLA9-dependent glucomannan has very few of these α-Gal branches, and that the hemicellulose is not distinguishable from AcGGM reported from other plants.

**Figure 2.**
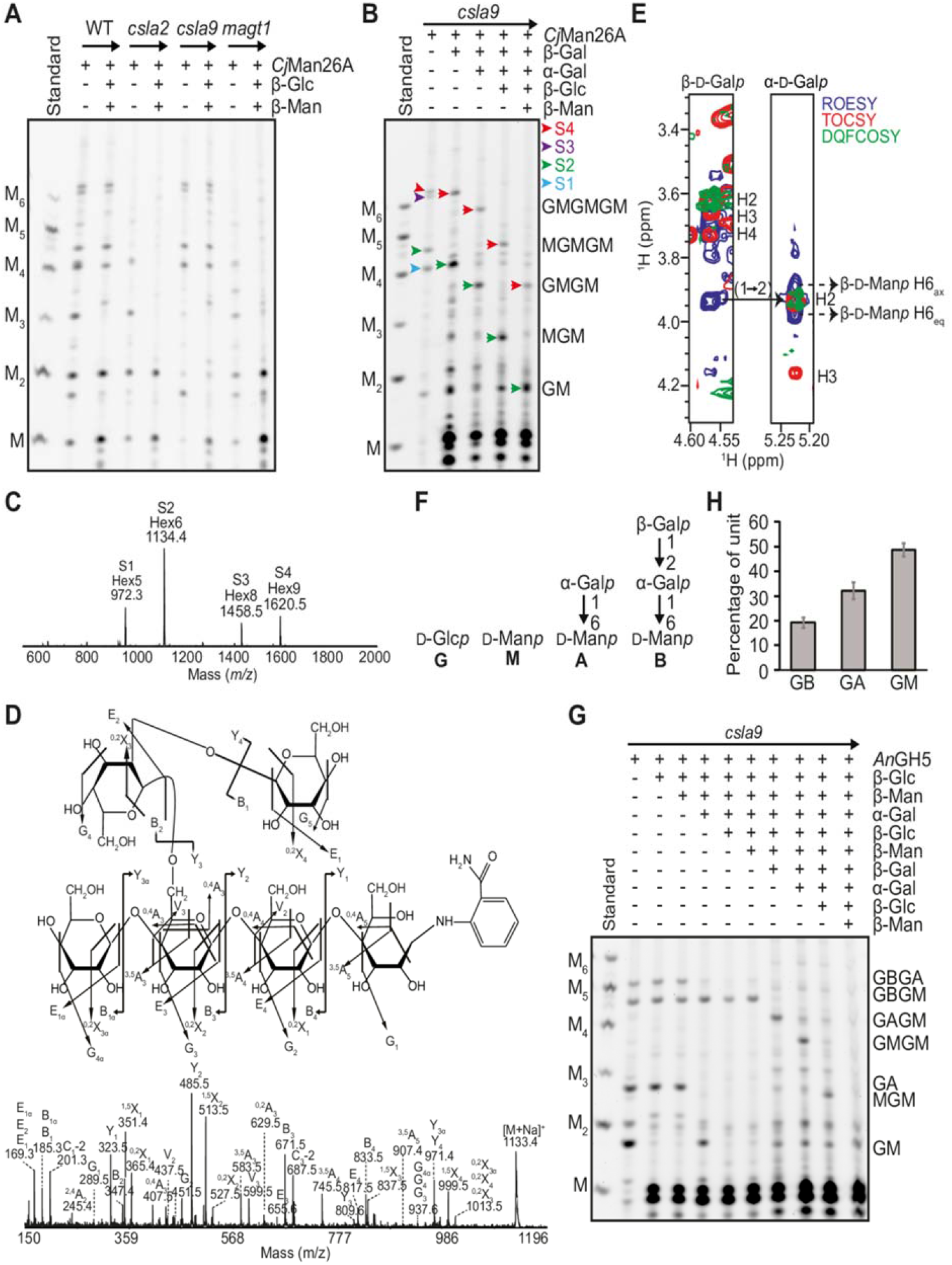
Structural analysis of β-galactosylated glucomannan oligosaccharides from Arabidopsis young stem. A, Characterization of glucomannan oligosaccharides released from WT, *csla2, csla9* and *magt1* cell walls by *Cj*Man26A. Glucomannan from *csla2* is degraded into M, MM, GMM and oligosaccharides migrating near M_4_. Many WT and *csla9* glucomannan oligosaccharides are resistant to β-glucosidase (β-Glc) and β-mannosidase (β-Man) enzyme digestions, whereas oligosaccharides from *csla2* are reduced to mono and disaccharides. B, Degradation of β-galactosylated glucomannan oligosaccharides from *csla9* young stem analysed by PACE. β-galactosidase (β-Gal), α-galactosidase (α-Gal), β-Glc, and β-Man enzymes were used sequentially. C, Products of *Cj*Man26A digestion of *csla9* cell walls were labelled with 2-AB and analysed by MALDI-TOF MS. The four main peaks correspond to the saccharides S1 to S4. D, S2 Hex6 in C was analysed by high-energy collision-induced dissociation (CID) MS/MS. The CID spectrum indicates that the α-Gal residue is linked to C-6 of the third hexose from the reducing end and that the β-Gal residue is linked to the C-2 or C-3 of the α-Gal. E, Nuclear magnetic resonance (NMR) analysis of S2. H-1 strip plots from 2D ^1^H-^1^H TOSCY (red), ROESY (blue), and DQFCOSY (green) spectra, showing the nuclear Overhauser effect (NOE) connectivity arising from the β-Gal*p*-1,2-α-Gal*p* linkage. F, A single-letter nomenclature for the identified β-GGM backbone and possible side chains. G, Characterization of *An*GH5 β-GGM glucomannan digestion products by PACE. *An*GH5 cleaves β-GGM from *csla9* young stem cell walls into GM, GA, GBGM, and GBGA oligosaccharides. H, Proportion of β-GGM disaccharides with different side chains from *An*GH5 digestion of etiolated *csla9* seedling glucomannan and PACE densitometry (*n* = 4). Error bars show the SD. Manno-oligosaccharide standards M to M_6_ are shown.

Next, we analysed the structure of CSLA2-dependent oligosaccharides released from *csla9* plants. We recently showed that the CSLA2-synthesised glucomannan in seed mucilage has a strictly repeating [4-Glc-β-1,4-Man-β-1,] disaccharide backbone with most of the Man residues substituted with α-1,6-Gal by the MAGT1 glycosyltransferase (Yu et al., 2018). Accordingly, to investigate if the oligosaccharides from etiolated seedlings were also α-galactosylated by MAGT1, we performed *Cj*Man26A digestions of *magt1* mutant seedling walls. The CSLA2-dependent oligosaccharides S1–S4 were absent or reduced in this mutant in all tissues, and two oligosaccharides corresponding to Glc-β-1,4-Man (GM) and Glc-β-1,4-Man-β-1,4-Glc-β-1,4-Man (GMGM) became more prominent (Figure 1 and Figure 2A). Therefore, CSLA2 likely synthesises a glucomannan with a repeating GM disaccharide backbone that is α-galactosylated by MAGT1.

To study the side chain structures in more detail, the four oligosaccharides S1–S4 were subjected to a sequential glycosidase digestion (Figure 2B). Since the presence of S1 to S4 is dependent on MAGT1, we investigated whether they are sensitive to α-galactosidase treatment. Interestingly, only the mobilities of S1 and S3, but not S2 and S4, were altered by α-galactosidase (Supplemental Figure S3A). The two α-galactosidase-treated oligosaccharides could be fully hydrolysed with alternating sequential β-glucosidase and β-mannosidase treatment, indicating that they are likely GMGM and GMGMGM. We analysed all four oligosaccharides S1-S4 by matrix-assisted laser desorption/ionization time-of-flight (MALDI-ToF) mass spectrometry (MS). The resultant spectra presented four main ions corresponding to S1 to S4, with mass Hex5 (m*/z* 972.3 [M+Na]^+^), Hex6 (*m/z* 1134.4 [M+Na]^+^), Hex8 (*m/z* 1458.5 [M+Na]^+^), and Hex9 (*m/z* 1620.5 [M+Na]^+^) respectively (Figure 2C). We reasoned that the mass of S1 and S3 likely correspond to Hex5 and Hex8, and that they carry one and two α-Gal residues, respectively. Subsequent analysis of the S1 ion by collision-induced dissociation (CID) MS/MS located its α-Gal branch to the first Man residue from the non-reducing end in the GMGM structure (Supplemental Figure S3B). Combined with the fact that *Cj*Man26A requires an unsubstituted Man at the −1 subsite for hydrolysis, these PACE and MS results indicate that all Man residues in S1 and S3 except the reducing end are α-galactosylated.

Oligosaccharides S2 and S4 were resistant to all the above glycosidase treatments (Supplemental Figure S3A), suggesting that these oligosaccharides had additional terminal substitutions. In tobacco cell cultures and kiwifruit, a glucomannan with β-1,2-Gal decorations on its α-1,6-Gal residues has been identified (Sims et al., 1997; Schröder et al., 2001). Interestingly, after β-galactosidase treatment, S2 and S4 co-migrated with S1 and S3 (Figure 2B). Sequential digestion with α-galactosidase, β-glucosidase and β-mannosidase confirmed that the β-galactosidase products had the same structure as S1 and S3. This indicates that S2 and S4 are S1 and S3 substituted with a β-Gal residue. Furthermore, CID MS/MS analysis of S2 showed that the second hexose from the reducing end is decorated with a hexose, which is itself substituted with a hexose, consistent with a β-Gal-α-Gal-disaccharide substitution of a backbone Man residue (Figure 2D). To confirm the linkage between the β-Gal and α-Gal, the S2 oligosaccharide was purified and analysed by 2D NMR. ^1^H and ^13^C chemical-shift assignments are shown in Supplemental Table S1. The β-Gal residue was deduced to link to the α-Gal residue *via* a 1,2-linkage due to the downfield shift of the α-Gal C-2 and an intense ROE peak between β-Gal H-1 and α-Gal H-2 (Figure 2E). Therefore, CSLA2 synthesizes a glucomannan with a repeating GM disaccharide backbone, on which the Man residues may be decorated with either single α-1,6-Gal or a β-1,2-Gal-α-1,6-Gal disaccharide.

We named this novel glucomannan β-GalactoGlucoMannan (β-GGM) because the β-Gal is one of the distinguishing features. By analogy to the XyG naming system, a one-letter code nomenclature was adopted to simplify the depiction of the arrangement of sugars and side chains along the backbone (Figure 2F). The letters G and M represent unsubstituted Glc and Man residues respectively. α-1,6-galactosylated Man residues are denoted by the letter A and the Man residues substituted by a Gal-β-1,2-Gal-α-1,6-disaccharide are denoted by the letter B. Using *An*GH5, which is a mannanase that can cleave following M or A units in a galactoglucomannan backbone (von Freiesleben et al., 2016). Digestion of β-GGM from the *csla9* young stem released four oligosaccharides (Figure 2G): GM, GA, GBGM, and GBGA. From these data, about 50% of backbone Man residues were decorated with α-1,6-Gal and about 40% of these α-Gal residues are further decorated with β-1,2-Gal (Figure 2H). Oligosaccharides with consecutive β-galactosylated Man residues were not seen (e.g. no GBGBGM, but GBGAGM and GBGM were seen), indicating that β-galactosylation is not random, but spaced at least four residues apart. Thus, in addition to the disaccharide backbone GM repeat, the β-GGM has a larger scale even-length pattern of at least four residues.

### β-GGM is widely present in eudicots

We considered whether β-GGM might be widespread in plants. As mentioned above, oligosaccharides that could arise from β-GGM were previously identified in tobacco cell cultures and kiwifruit samples (Sims et al., 1997; Schröder et al., 2001). We performed *An*GH5 mannanase digestions on alkali-extracted mannan from PCW-rich samples from tomato fruits, kiwi fruits and apple fruits, representatives from the asterid and rosid eudicot clades. The β-GGM representative oligosaccharides GBGM and GAGM were present (Supplemental Figure S4, A to C). GBGM was digested by the β-galactosidase. Tomato fruit showed a higher proportion of GBGM oligosaccharide than the other plant tissues (Supplemental Figure S4), indicating that there is some variability in the level of β-Gal substitution of β-GGM.

We note that Arabidopsis seed mucilage glucomannan has a structure similar to β-GGM except that the β-Gal substitution was not reported (Yu et al., 2018). The MUM2 β-galactosidase is highly expressed in seed mucilage and has been shown to remove pectin terminal β-Gal (Dean et al., 2007; Macquet et al., 2007). We therefore considered the possibility that MUM2 also acts on β-GGM in mucilage to remove any β-Gal decoration. To test this hypothesis, alkali-extracted *mum2* mucilage was treated with *Cj*Man26A and analysed by PACE. Minor GBGM and GBGAGM oligosaccharides were clearly present (Supplemental Figure S4D). We therefore conclude that mucilage mannan is also β-GGM, and it has been partly trimmed by the MUM2 β-galactosidase. Arabidopsis mucilage glucomannan is not as unusual as previously thought (Yu et al., 2018), but another example of a tissue with β-GGM.

### AT4G13990 from GT47 clade A encodes β-GGM β-galactosyltransferase, MBGT1

To understand the β-GGM biosynthesis, we attempted to identify the Mannan β-GalactosylTransferase (MBGT). We noted that β-GGM and XyG share high structural and biosynthetic similarities, summarised here and in Figure 3A. Both backbones have β-1,4-Glc residues in the backbone, which in β-GGM alternate with β-1,4-Man residues (Man differs from Glc only in epimerisation of the C-2 OH). Both backbones are made by closely related GT2 members: the XyG backbone is synthesized by CSLCs (Cocuron et al., 2007; Liepman et al., 2007; Kim et al., 2020), while the β-GGM backbone is synthesized by a CSLA. Furthermore, the first side chain sugars are attached to the C-6 OH of Glc on the XyG backbone and to the C-6 OH of Man on the β-GGM backbone. The XyG α-1,6-Xyl is transferred by XXTs and α-1,6-Gal is transferred to β-GGM by MAGT1, both from the GT34 family (Scheller and Ulvskov, 2010). The disaccharide branch second sugar in β-GGM is β-1,2-Gal. The same sugar and linkage is found in XyG.

**Figure 3.**
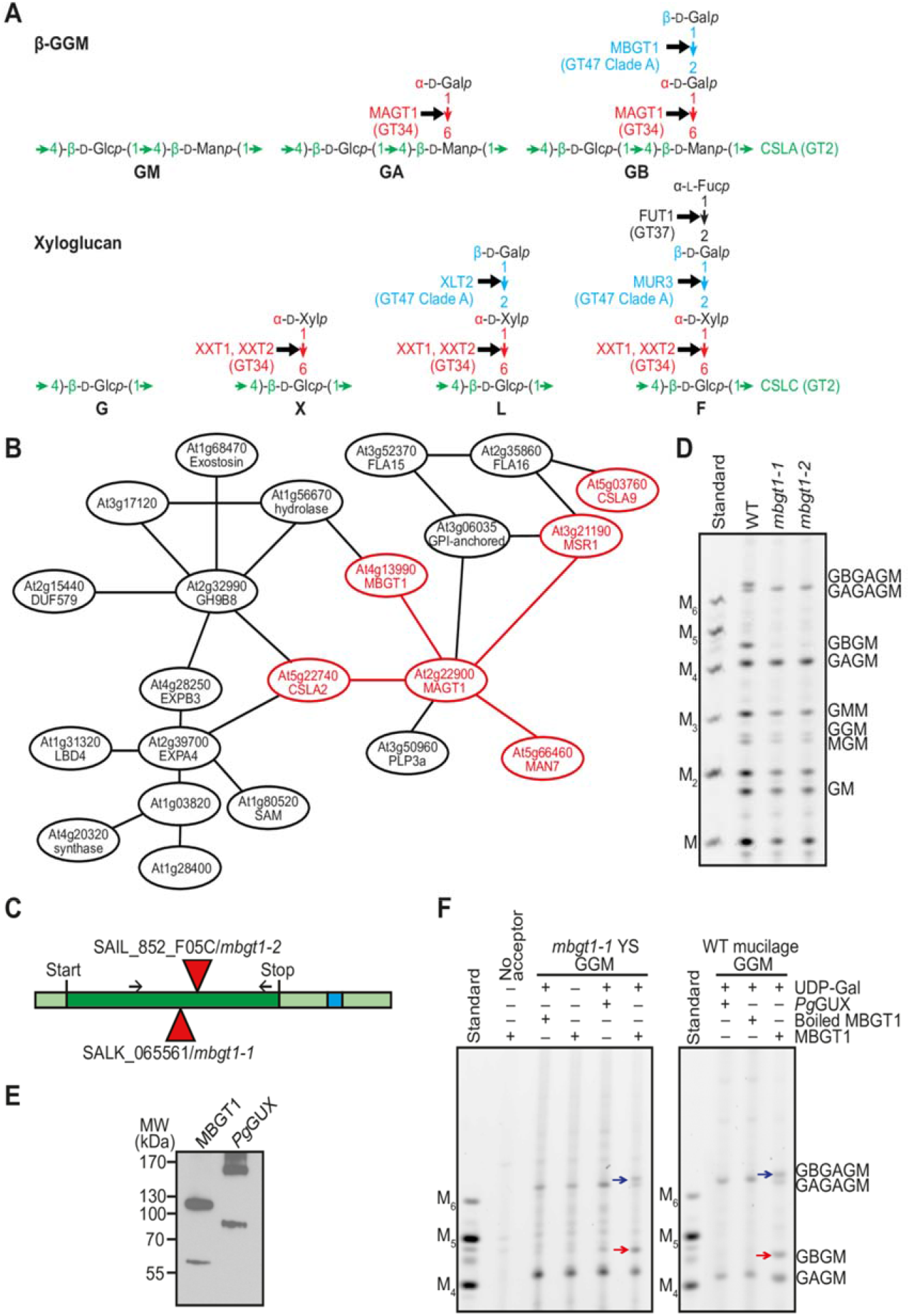
AT4G13990 from CAZy GT47 Clade A encodes Arabidopsis mannan β-galactosyltransferase 1 (MBGT1). A, β-GGM and XyG share structural and biosynthesis similarities. These two polysaccharides exhibit analogous linkages in their backbones and corresponding side chain sugars. For position in the hemicellulose, the responsible glycosyltransferases are from the same CAZy family. B, AT4G13990/MBGT1 from GT47 Clade A is in a co-expression network with CSLA2 and other mannan-related genes. C, Gene model representing *MBGT1*. Red triangles represent the position of T-DNA insertions in mutant lines analysed in this study. Dark green represents the exon. Light green represents the UTR and blue shows an intron. D, Stem material of two insertional mutants of the *MBGT1* gene was analysed by PACE by *Cj*Man26A. No β-galactosylated oligosaccharide was detected in either *mbgt1* mutant. E, Western blot of 3×Myc-tagged recombinant proteins expressed in *N. benthamiana*. The expected mass of 3×Myc–MBGT1 is 64.86 kDa. The expected mass of the control enzyme 3×Myc–*Pg*GUX is 78.18 kDa. Both proteins form stable dimers. F, *In vitro* activity of the recombinant MBGT1 protein. In the left panel, *mbgt1-1* young stem (YS) glucomannan was used as an acceptor for MBGT1-mediated galactosylation, whereas in the right panel, WT adherent mucilage glucomannan was used. The products were analysed with PACE using digestion with *Cj*Man26A. Arrows indicate band shifts after each reaction. Manno-oligosaccharide standards M to M_6_ are shown.

Given these extensive XyG and β-GGM similarities, we hypothesised that MBGT might be found in GT47 clade A, which contains many XyG β-glycosyltransferases (MUR3, XLT2, and XUT1) (Geshi et al., 2018) and also many putative GTs with no known functions (classified *At*GT11-*At*GT20 in (Li et al., 2004)). To identify MBGT candidates, we constructed a comprehensive phylogeny of GT47-A sequences from across the plant kingdom. We collected GT47-A sequences from the genomes of 96 streptophytes (listed in Supplemental Table S2) and inferred an unrooted phylogeny (Figure 4). The sequences were clustered into at least seven groups: group I (containing only non-spermatophyte sequences, but including previously characterised *Pp*XLT2 and *Pp*XDT enzymes from *Physcomitrium patens* (Zhu et al., 2018)), group II (containing *At*XUT1 (Pena et al., 2012) and *At*GT20), group III (*At*XLT2 (Jensen et al., 2012), *Os*XLT2 from rice (Liu et al., 2015), and tomato enzymes *Sl*XST1 and *Sl*XST2 (Schultink et al., 2013)), group IV (*At*GT19), group V (*At*GT17), group VI (*At*MUR3 (Madson et al., 2003), *Sl*MUR3 (Schultink et al., 2013), and *Os*MUR3 (Liu et al., 2015)), and group VII (*At*GT11–15). Because none of the enzymes in groups IV, V and VII had been characterised, we considered these groups to be a potential source of new activities (although *At*GT11 has recently been implicated in XyG synthesis in pollen tubes (Wei et al., 2021)). Accordingly, for each Arabidopsis gene within these GT47-A groups, we analysed its co-expression using the co-expression database tool ATTED-II (Obayashi et al., 2018). Interestingly, we found that *At4g13990* (*At*GT14, group VII) is co-expressed with the glucomannan biosynthetic enzymes CSLA2, MAGT1, and MSR1 (Figure 3B). Hence, we considered the possibility that *At4g13990* could encode MBGT.

**Figure 4.**
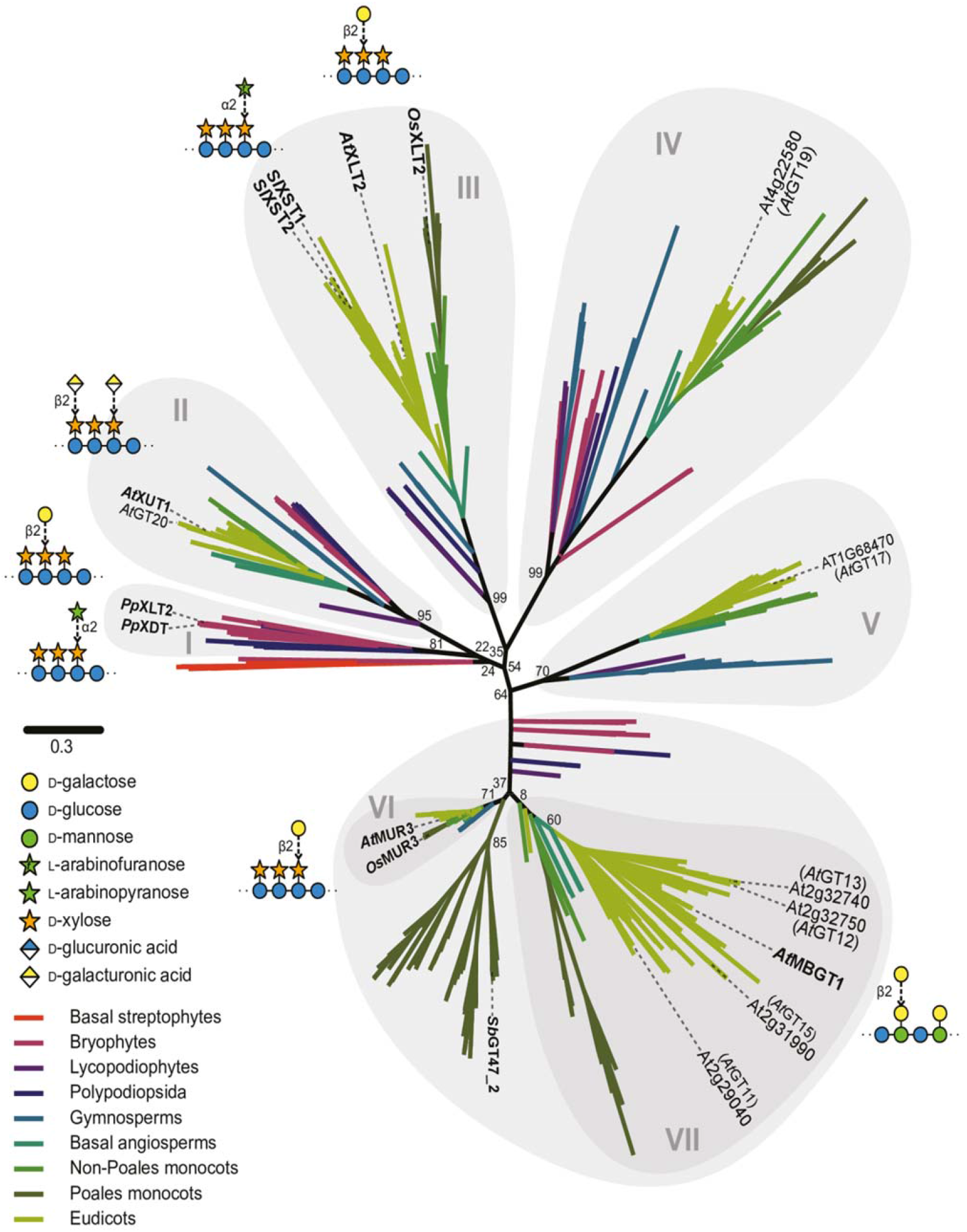
Un-rooted phylogenetic tree of CAZy GT47 Clade A. Sequences from the genomes of 96 streptophytes (Supplementary Table 2) were used to construct a comprehensive phylogeny of GT47 Clade A. Most sequences were downloaded from PLAZA (https://bioinformatics.psb.ugent.be/plaza/), but were supplemented with additional sequences from further genomes, derived from HMMER and TBLASTN searches. The basal streptophyte representative is *Klebsormidium nitens*, and the Lycopodiophyte representative is *Selaginella moellendorffii*. Sequences were aligned with MAFFT and truncated to leave only the predicted GT47 domain. The phylogeny was then inferred using FastTree, with 100 bootstrap pseudo-replicates. Percentage replication is indicated for important splits. The resultant tree revealed the existence of seven main subgroups within GT47-A (groups I–VII), four of which contain known XyG glycosyltransferases. The group containing MBGT was designated group VII. The activities of characterised enzymes are illustrated in SNFG format.

To assess the potential role of At4g13990/*At*GT14 in β-galactosylation of β-GGM, cell walls from young stems of two homozygous knockout *At4g13990* lines (named *mbgt1-1* and *mbgt1-2*, Figure 3C) were digested with *Cj*Man26A, and the products were analysed by PACE. Remarkably, both mutant lines lacked the β-galactosylated β-GGM oligosaccharides (Figure 3D), indicating that this enzyme is required for normal β-galactosylation of β-GGM.

To confirm the activity of At4g13990/*At*GT14, we conducted an assay for MBGT activity *in vitro* using AT4G13990 protein transiently expressed in tobacco leaves. Alkali-treated cell wall materials from *mbgt1-1* young stem and WT adherent mucilage, rich in β-GGM but lacking the β-galactosylation (Yu et al., 2018), were used as acceptors. To detect β-galactosylated glucomannan, the assay products were digested with mannanase *Cj*Man26A and analysed by PACE. In the presence of UDP-Gal and microsomes from tobacco expressing At4g13990/*At*GT14, β-GGM oligosaccharides were produced from mucilage and young stem acceptors (Figure 3, E and F). In contrast, when microsomes from tobacco over-expressing *Picea glauca* GlucUronic acid substitution of Xylan (*Pg*GUX1) (Lyczakowski et al., 2017) were used as the control enzyme, no β-galactosylation was detected. Taken together with the mutant plants, these results confirm that At4g13990/*At*GT14 encodes MBGT, and so we named it MBGT1.

### Arabidopsis mutants in β-GGM and XyG side chain structure show negative genetic interactions

The structural and biosynthetic relationships between β-GGM and XyG suggest that these two polysaccharides may play related functions *in vivo*. If our hypothesis is correct, β-GGM biosynthesis disruption might exacerbate the phenotypes of XyG synthesis mutants.

Mutant plants lacking β-GGM β-Gal (*mbgt1-1*) grew indistinguishably from wild type plants (Figure 5). An analogous mutant in XyG is *mur3-3*, which lacks the third position β-Gal. It has a cabbage-like growth phenotype with curled rosette leaves, and short stems (Tamura et al., 2005; Tedman-Jones et al., 2008). We generated *mbgt1-1 mur3-3* double mutant plants. As expected, they had no detectable β-GGM with B units and XyG with no third position L and F (β-Gal further substituted with Fuc) units (Supplemental Figure S5, A and B). Interestingly, these β-GGM and XyG double mutants had a smaller rosette than *mur3-3*, with more severely curled rosette leaves (Figure 5A). In addition, the inflorescence stem was shorter than the *mur3-3* single mutant plants (Figure 5 C). The allelic Arabidopsis *mur3-1* mutant, with a single-point mutation in MUR3 (Madson et al., 2003; Jensen et al., 2012) also has defective XyG. For unclear reasons, this XyG mutant does not exhibit a cabbage phenotype, but the plants are shorter and has an increased number of rosette and cauline branches than WT (Jensen et al., 2012). To test for genetic interactions with this allele, we generated *mbgt1-1 mur3-1* double mutant plants. Compared to the single mutant *mur3-1* plants, the *mbgt1-1 mur3-1* double mutant was significantly shorter and had more cauline branches (Figure 5, B-F). The increased severity of the *mur3-1* phenotypes when combined with *mbgt1-1* indicates that β-galactosylation of β-GMM is important for β-GMM function, and suggests that the disaccharide side chains in both polysaccharides have similar functions.

**Figure 5.**
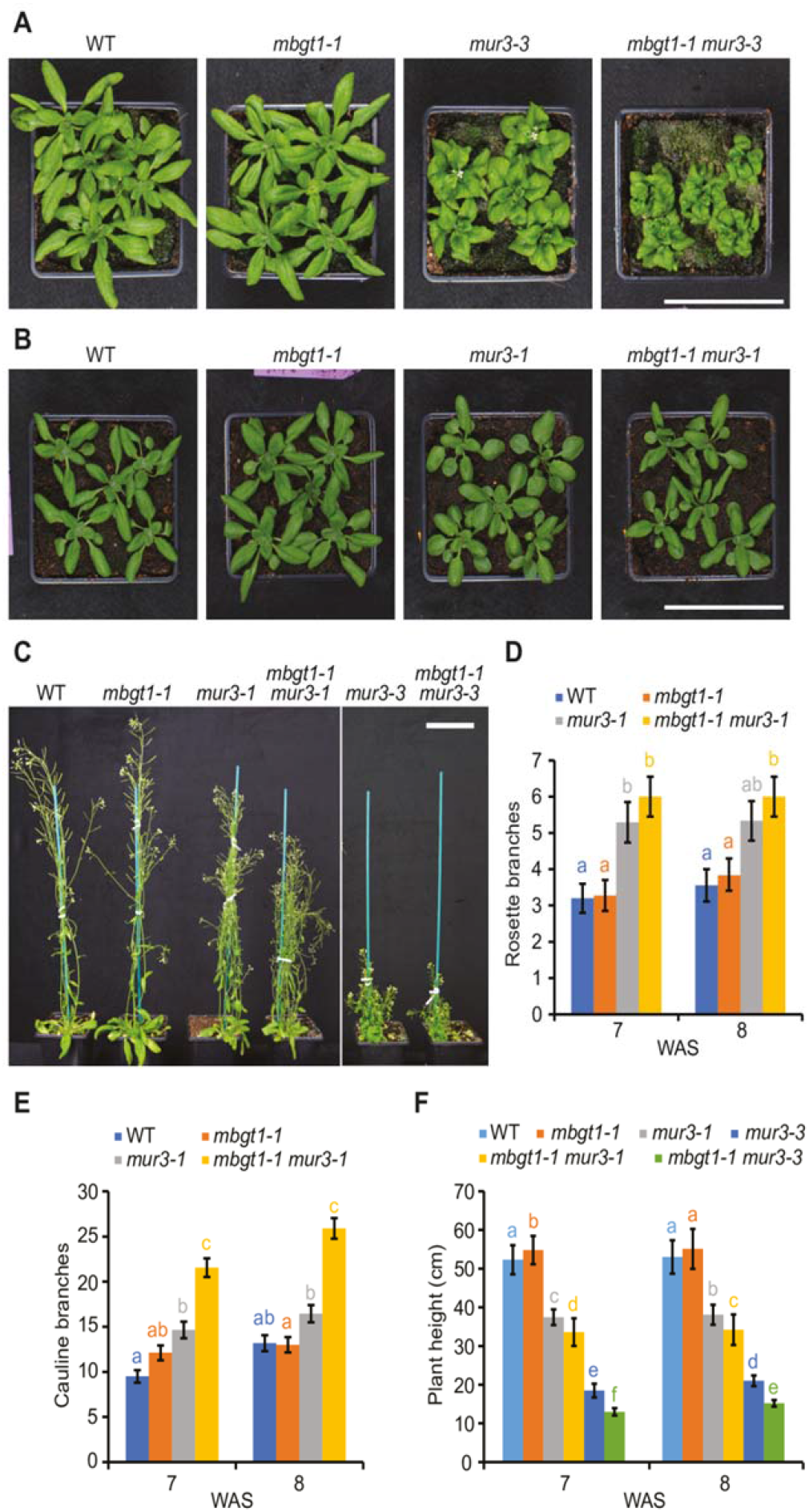
The importance of β-galactosylation of β-GGM is revealed in the XyG β-galactosylation mutant *mur3*. A, Four-week-old rosettes of *mur3-3* T-DNA insertion mutant and *mbgt1-1 mur3-3* double mutant. B, Four-week-old rosettes of *mur3-1* point mutant and *mbgt1-1 mur3-1* double mutant. C, Six-week-old plants, showing dwarfing of the *mur3* and *mbgt1-1 mur3* double mutants. D, E Quantification of the number of rosette branches (D) and cauline branches (E) for 7 and 8-week-old *mur3-1* and *mbgt1-1 mur3-1* plants. *mbgt1-1 mur3-1* mutants show no significant change in rosette branches, but a significant increase in cauline branches compared to *mur3-1*. Data were modelled by Poisson regression; a likelihood ratio test indicated a significant contribution of genotype in determining the number of stems (Rosette branches 7 weeks: *n* = 75, *G*^2^_3_ = 26.2, *p* = 8.6 × 10^−6^; 8 weeks: *n* = 74, *G*^2^_3_ = 16.6, *p* = 8.4 × 10^−4^). Cauline branches 7 weeks: *n* = 75, *G*^2^_3_ = 109, *p* < 2.2 × 10^−16^; 8 weeks: *n* = 73, *G*^2^_3_ = 144, *p* = 1.5 × 10^−24^). Results of post-hoc pairwise comparisons (within each time point) are indicated by compact letter display (letter sharing indicates lack of significant difference *i*.*e*. where *p* > 0.05). Data were modelled by Poisson regression; a likelihood ratio test indicated a significant contribution of genotype in determining the number of stems (Rosette branches 7 weeks: *n* = 75, *G*^2^_3_ = 26.2, *p* = 8.6 × 10^−6^; 8 weeks: *n* = 74, *G*^2^_3_ = 16.6, *p* = 8.4 × 10^−4^). Cauline branches 7 weeks: *n* = 75, *G*^2^_3_ = 109, *p* < 2.2 × 10^−16^; 8 weeks: *n* = 73, *G*^2^_3_ = 144, *p* = 1.5 × 10^−24^). Error bars represent standard error of the mean. F, Quantification of plant height for 7 and 8-week-old plants. One-way ANOVA indicated a significant contribution of genotype in determining plant height at both timepoints (7 weeks: *n* = 208, *F*_5,202_ = 1257, *p* < 2 × 10^−16^; 8 weeks: *n* = 200, *F*_5,194_ = 760, *p* < 2 × 10^−16^). Results of post-hoc pairwise comparisons (within each time point) are indicated by compact letter display. Apart from the significant difference between WT and *mbgt1-*1 at 7 weeks, where *p* = 0.0066, *p* < 1 × 10^−6^ for all significant differences. Error bars represent standard deviation. WAS, week after sowing. Scale bars = 9 cm.

### Arabidopsis mutants lacking β-GGM and XyG show negative genetic interactions

The *xxt1 xxt2* mutant, lacking detectable amounts of XyG, exhibits some morphological phenotypes in many tissues, yet the plants are able to grow relatively normal. To investigate if the absence of β-GGM exacerbates the phenotype of these plants, we crossed the *csla2* mutant with *xxt1 xxt2*. As previously reported, compared to wild type, the *xxt1 xxt2* mutant had narrow leaves and a smaller rosette diameter, somewhat shorter plants at 8 weeks, and shorter siliques (Kong et al., 2015) (Figure 6). The *csla2* mutant plants, lacking β-GGM, grew normally. Interestingly, the *csla2 xxt1 xxt2* mutant plants, lacking both β-GGM and XyG (Supplementary Figure 5C), had a more severe phenotype with slightly changed rosette appearance, significantly shorter stems at 6 to 8 weeks and shorter siliques than *xxt1 xxt2* (Figure 6).

**Figure 6.**
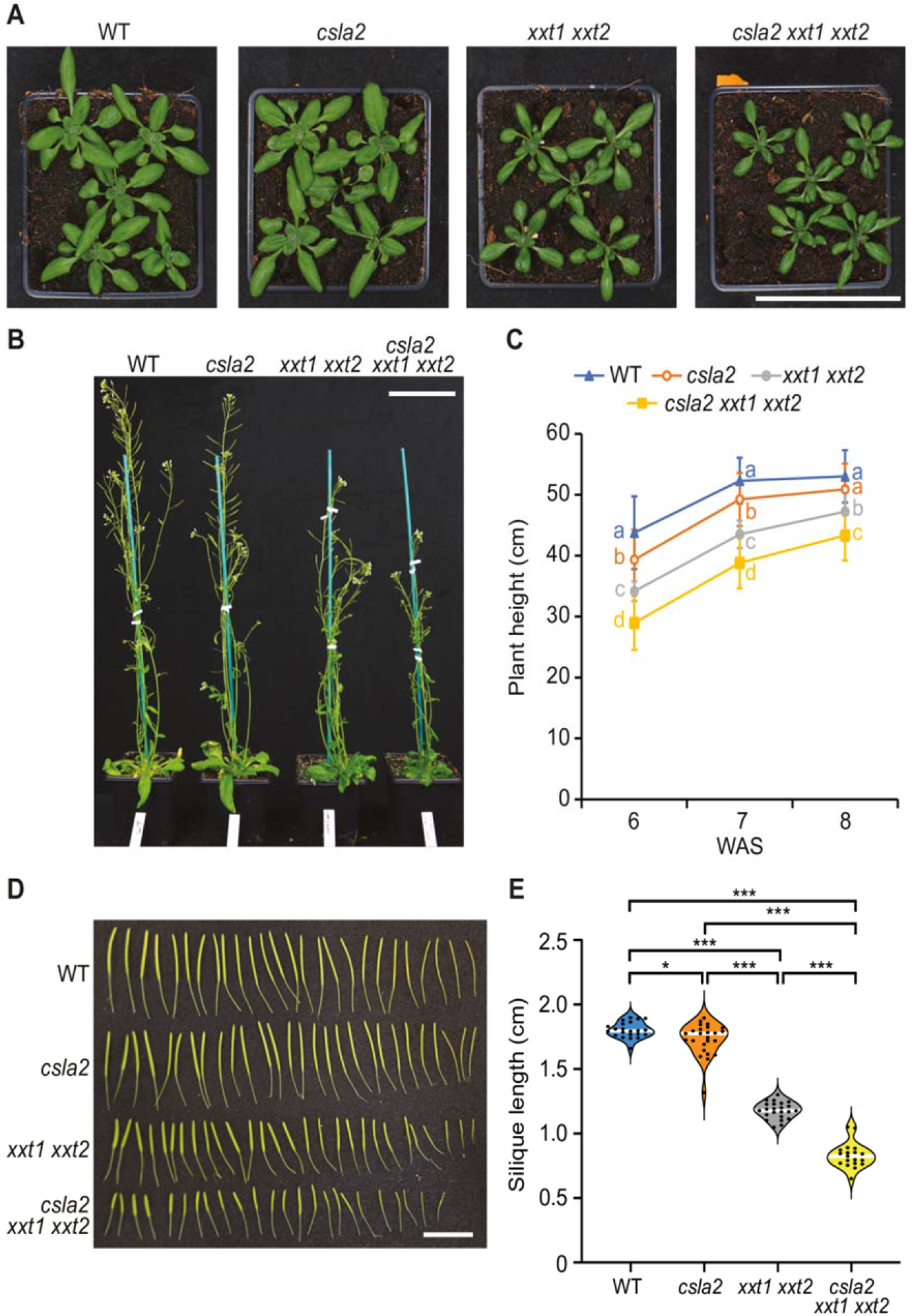
β-GGM function in primary cell walls is revealed when the XyG is missing. A, Four-week-old rosettes. Scale bar = 9 cm. B, Six-week-old plants. Scale bar = 9 cm. C, Quantification of plant height for 6, 7, and 8-week-old plants. One-way ANOVA indicated a significant contribution of genotype in determining plant height at all three timepoints (6 weeks: *n* = 131, *F*_3,127_ = 65.0, *p* < 2 × 10^−16^; 7 weeks: *n* = 136, *F*_3,132_ = 88.2, *p* < 2 × 10^−16^; 8 weeks: *n* = 131, *F*_3,127_ = 35.8, *p* < 2 × 10^−16^). Results of post-hoc pairwise comparisons (Tukey’s honest significant difference) are indicated by compact letter display. For all significant differences, *p* < 0.001 apart from WT–*clsa2* at week 7 (*p* = 0.0063) and *csla2*–*xxt1 xxt2* at week 8 (*p* = 0.0026). D, Siliques from 7-week-old plants. Scale bar = 2 cm. E, Violin plot of silique length. Siliques from more than three plants were measured for each genotype. Black circles indicate individual measurements; white lines represent the group mean, and error bars indicate standard deviation. One-way ANOVA indicated a significant contribution of genotype in determining plant height at all three time points (*n* = 89, *F*_3,85_ = 553, *p* < 2 × 10^−16^). Results of post-hoc pairwise comparisons (Tukey’s honest significant difference; WT, *n* = 22; *csla2, n* = 25; *xxt1 xxt2, n* = 23; *csla2 xxt1 xxt2, n* = 19) are indicated with asterisks.

Next, we investigated if the *xxt1 xxt2* phenotype in etiolated hypocotyls was affected by the loss of β-GGM. Plant lines were grown on MS plates in the dark for between 3 and 7 days to measure hypocotyl length. Hypocotyl length differences between the mutants became evident 4 days after germination. Up to day 7, no significant difference was observed between *csla2* and WT seedlings. *xxt1 xxt2* seedlings were shorter than those of WT, consistent with previously published results (Xiao et al., 2016). *csla2 xxt1 xxt2* etiolated seedlings exhibited even shorter hypocotyls than those of *xxt1 xxt2* (Figure 7, A and B). In addition, the *csla2 xxt1 xxt2* seedlings have perturbed growth showing some twisting of hypocotyls (Figure 7A). These results suggest that β-GGM and XyG have connected functions in normal plant development.

**Figure 7.**
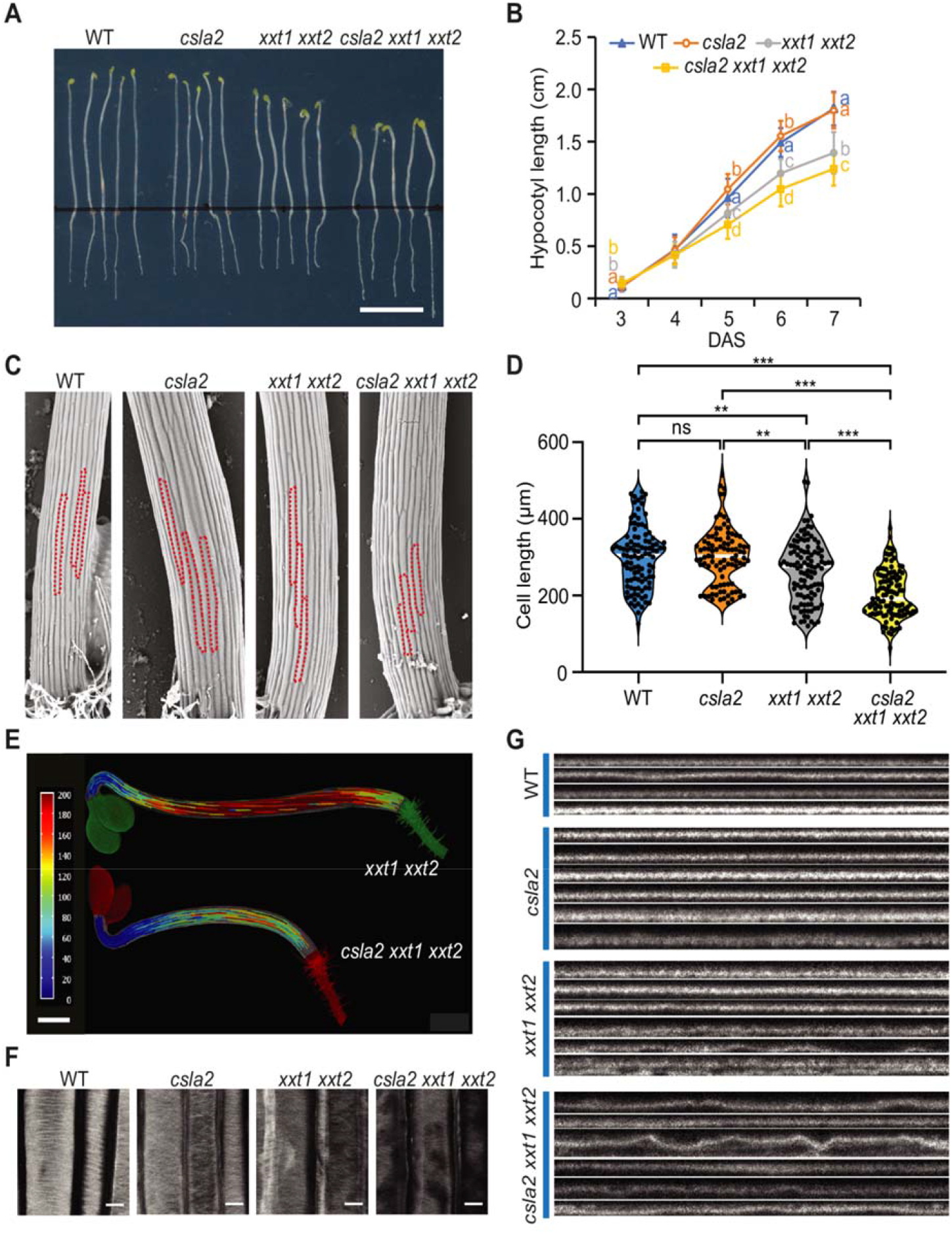
A role of β-GGM in cell expansion and cellulose organisation. A, Six-day-old hypocotyls grown on MS medium with sucrose. Scale bar = 1 cm. B, Quantification of hypocotyl length for 3- to 6-day-old seedlings (n ≥ 40 seedlings for each point per genotype). DAS, days after sowing. Error bars represent standard deviation. Although one-way ANOVA indicated no significant difference between genotypes at 4 days (*n* = 213, *F*_3,209_ = 2.58, *p* = 0.054), a significant difference was seen at 3 days, 5 days and after (3 days: *n* = 197, *F*_3,193_ = 40.7, *p* < 2 × 10^−16^; 5 days: *n* = 276, *F*_3,272_ = 82.8, *p* < 2 × 10^−16^; 6 days: *n* = 271, *F*_3,267_ = 177, *p* < 2 × 10^−16^; 7 days: *n* = 245, *F*_3,241_ = 167, *p* < 2 × 10^−16^). Results of post-hoc pairwise comparisons (Tukey’s honest significant difference) are indicated by compact letter display. C, Cryo-SEM analysis of 4-day-old etiolated hypocotyls from WT and mutant plants. Individual cells in the tissue are outlined. Cells are shorter in the *csla2 xxt1 xxt2* triple mutant than in the *xxt1 xxt2* double mutant. Scale Bar = 100 μm. D, Quantification of cell length of 4-day-old hypocotyls. Black circles indicate individual measurements; white lines represent the group mean One-way ANOVA indicated a significant contribution of genotype in determining hypocotyl cell length (*n* = 413, *F*_3,409_ = 40.44, *p* < 2 × 10^−16^). Results of post-hoc pairwise comparisons (Tukey’s honest significant difference) are indicated by asterisks (* *p* < 0.05, ** *p* < 0.01, *** *p* < 0.001). E, Heatmap showing 4-day-old hypocotyl cell length. Scale Bar = 500 μm. F, G Four-day-old hypocotyls were stained with Pontamine S4B and then observed under a confocal microscope. Representative image of hypocotyl primary cell wall (F). A survey of orthogonal views showing the profile of the hypocotyl primary cell wall (G). Scale bars = 10 μm.

### Defects in cell elongation and cellulose microfibril organisation

To investigate the developmental changes in β-GGM and XyG mutants, we imaged four-day-old etiolated hypocotyls by cryo-SEM and studied the epidermal cell lengths. Compared to WT, the *xxt1 xxt2* mutant exhibited a small reduction in cell length, while the *csla2 xxt1 xxt2* exhibits a larger reduction (Figure 7, C and D). To better visualize the differences in cell expansion along the hypocotyl, we imaged and computationally segmented the hypocotyl cells of the *xxt1 xxt2* and *csla2 xxt1 xxt2* mutants. A heat map of cell length demonstrates cells are consistently shorter in the *csla2 xxt1 xxt2* mutant along the whole hypocotyl (Figure 7E). The data suggest cell expansion is further reduced, compared to the loss of XyG alone, by the absence of both β-GGM and XyG.

XyG mutants have revealed the importance of the polysaccharide for normal cellulose fibril arrangements and wall formation (Xiao et al., 2016; Kim et al., 2020). We hypothesised that loss of β-GGM and XyG may affect cell elongation by altering cellulose microfibril arrangements. We processed and stained the cellulose using pontamine fast scarlet 4B dye (Thomas et al., 2017) and imaged epidermal cells using confocal microscopy (Figure 7F). Stained transverse bundles could be observed for WT and *csla2* and these bundles were less defined with areas of missing signal in the *xxt1 xxt2* and *csla2 xxt1 xxt2* mutants suggesting uneven walls. Orthogonal profiles along the cell length and through the stained walls show thin and even walls for WT and *csla2*, but uneven, “rippled” profiles for *xxt1 xxt2*. This effect is worsened in *csla2 xxt1 xxt2* (Figure 7G). Although the features could be an effect of the processing steps of pontamine staining, they reveal differences in the cellulose arrangements in the cell walls of the mutants that are dependent on the presence of β-GGM and XyG.

### β-GGM has low mobility in primary cell walls

Hemicellulose polysaccharides that are bound to cellulose are relatively immobile in the cell wall (Bootten et al., 2004). Solid-state NMR (ssNMR) can be used to distinguish more mobile constituents from these relatively immobile polymers. For example, ^13^C cross-polarisation (CP)-magic-angle spinning (MAS) ssNMR has been used to study xyloglucan, xylan and glucomannan bound to cellulose. On the other hand, soluble polymers can be seen by direct polarisation (DP)-MAS ssNMR (Metz et al., 1994; Simmons et al., 2016; Cresswell et al., 2021). Because of the relatively low abundance of β-GGM in plants, and to study a simplified PCW, we exploited Arabidopsis callus cultures of hemicellulose biosynthesis mutants. Compared to seedlings, Arabidopsis callus cultures are relatively homogenous and reproducible between many genotypes. The cells synthesise polysaccharides typical of primary cell walls (Prime et al., 2000; Nikolovski et al., 2012) and can be labelled by growing with ^13^C-glucose. This enables two-dimensional spectra, in particular using the through-bond refocused INADEQUATE experiment, to be recorded. Such spectra of wild type callus cells are observed to be complex, and XyG signals dominate (Supplementary Figure S7). Thus, we generated *irx9l xxt1 xxt2* callus cultures to remove both XyG and xylan to simplify the spectra as much as possible, leaving β-GGM as the main hemicellulose. To help in assigning the β-GGM signals in the spectra, we also generated and analysed a *csla2 xxt1 xxt2* mutant callus, which lacks the β-GGM as well as XyG (Supplemental Figure S6, Supplementary Table S1). We carried out ssNMR on cell walls without drying or pretreatments, to preserve native arrangements of polymers as much as possible. Figure 8 shows that both Man residues and α-Gal branches of β-GGM can be seen in a ^13^C CP-INADEQUATE MAS NMR spectrum that detects relatively immobile polymers such as cellulose and bound hemicelluloses. The ^13^C NMR chemical shifts of these β-GGM residues are consistent with those from extracted Kiwi fruit glucomannan (Schröder et al., 2001) (Supplemental Table 1), supporting the assignments. β-Gal was not detected in the solid-state NMR spectra, perhaps due to lower frequency or higher mobility of this substitution. The similarity in ^13^C shifts to the previous solution-state assignments suggests that there are no major β-GGM conformational differences in the cell wall. The ^13^C shifts of these β-GGM residues are distinct from those of AcGGM (Terrett et al., 2019; Cresswell et al., 2021), consistent with the different chemical structure of these polymers. Importantly, like XyG and cellulose, β-GGM was not detected in DP-INADEQUATE spectra, in which mobile polymers are seen. Having assigned the β-GGM spectral peaks, we similarly investigated WT cell walls, and confirmed β-GGM is also detectable in CP-INADEQUATE spectra in the presence of the XyG and xylan (Supplementary Figure S7). Due to the low abundance of β-GGM, we have not been able to conduct through-space ssNMR experiments to investigate β-GGM proximity with cellulose. Nevertheless, the experiments indicate that β-GGM has limited mobility in the wall consistent with binding to cellulose.

**Figure 8.**
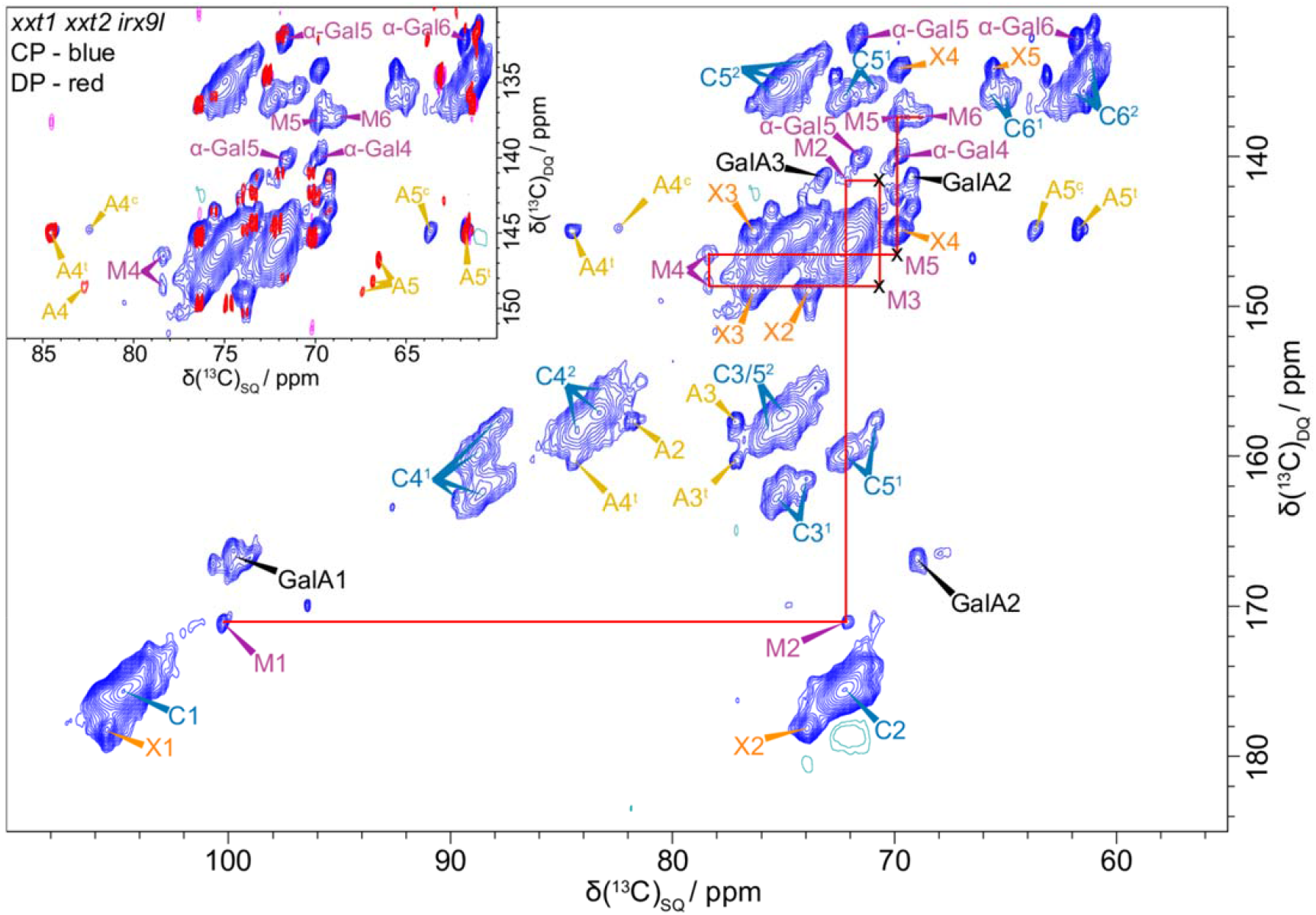
^13^C CP- and DP-refocused INADEQUATE MAS solid-state NMR spectra show the β-GGM peaks in *irx9l xxt1 xxt2* callus. The β-GGM peaks are labelled: mannose (M) and α-Gal. Also labelled are the main cellulose peaks (domain 1, C^1^; domain 2, C^2^), galacturonic acid (GalA) of pectin, and a terminal xylose (X) linked to an unknown polymer. A terminal arabinose (A^t^) and another arabinose (A^c^) are also labelled. The inset shows an overlay of the CP (blue) and DP (red) INADEQUATE spectra for the M5, M6 region. It is clear that M5 and M6 are not visible in the DP spectrum, i.e. are not mobile. Spectra were acquired at a ^13^C Larmor frequency of 251.6 MHz and a MAS frequency of 12.5 kHz. The spin-echo duration used was 2.24 ms.

## Discussion

XyG has been the focus of eudicot PCW hemicellulose functional studies because of its abundance, and it is the only eudicot hemicellulose with a clear role in cell wall elongation (Burton et al., 2010; Park and Cosgrove, 2015). Here, we report a widespread patterned glucomannan that shows structural and biosynthetic similarities to XyG, which we name β-GGM (Supplemental Figure S1). These two polysaccharides have related roles in cell elongation in plant development, with a role of β-GGM becoming more evident in tissues or mutants without functional XyG. Studies of the role of hemicelluloses in PCW architecture and function should now consider contributions by both polysaccharides.

The glucomannan and XyG biosynthetic enzymes are evolutionarily related (Yin et al., 2009; Wang et al., 2020). The β-GGM backbone is synthesized by CSLA2 in Arabidopsis, a CAZy family GT2 enzyme. Within plant GT2 enzymes, the CSLA enzymes are most closely related to the CSLC family (Mikkelsen et al., 2014; Wang et al., 2020), which are the XyG backbone synthases (Cocuron et al., 2007; Kim et al., 2020). In Arabidopsis, CSLA9 is required for biosynthesis of AcGGM, the random patterned, acetylated glucomannan in tissues with SCWs (Goubet et al., 2009). Therefore, there may be functional specialisation within the CSLA enzyme family. Whether the ability to make the patterned backbone for β-GGM is intrinsic to specific CSLA enzymes or induced by factors such as MSR proteins (Voiniciuc et al., 2019; Robert et al., 2021) remains to be investigated in plants. The land plant GT2 CSLAs have evolved from the streptophyte algal CSLA/K family (Wang et al., 2020) which is likely to synthesise a mannan. There is no report to our knowledge of glucomannans before the evolution of land plants, so this enzyme may synthesise a homomannan. The early land plants have been reported to have an acetylated glucomannan (Geddes and Wilkie, 1972; Popper and Fry, 2003; Nothnagel and Nothnagel, 2007; Zhang et al., 2014), suggesting that glucomannans are a land plant adaptation. The side chain biosynthesis of β-GGM and XyG is also related. The α-1,6-galactosyltransferase MAGT1 for β-GGM and α-1,6-xylosyltransferase XXTs for XyG are all from the GT34 families (Scheller and Ulvskov, 2010; Yu et al., 2018). We recently showed that MAGT1 activity has the ability to galactosylate Man in the patterned β-GGM backbone (Yu et al., 2018), but other MAGTs may show preferences for different Man or Glc residue arrangements. Here, we also identified the enzyme making the β-1,2-Gal disaccharide branch, MBGT1. It is from GT47 Clade A, which contains the XyG β-1,2-Gal transferases amongst many other XyG active enzymes. These extensive similarities in biosynthetic enzymes may imply that β-GGM and XyG have a common ancient evolutionary origin, for example in streptophyte algae where XyG and CSLA/K were present (Mikkelsen et al., 2014; Mikkelsen et al., 2021). In this scenario, both polysaccharides persisted through land plant evolution to modern eudicots. Alternatively, the β-GGM biosynthesis may have arisen during land plant evolution from the acetylated glucomannan biosynthesis pathway. We have not yet studied the presence of β-GGM across the plant kingdom, and so we are unable to determine yet whether the ability to make β-GGM is ancient or alternatively arose during land plant evolution. Evolution of the synthesis β-GGM would require divergence of CSLAs to make the patterned *vs* unpatterned backbones, specialisation of GT34s to add galactose to the patterned backbone, and alteration of a XyG GT47 activity for generation of the β-GGM disaccharide side chains. This second hypothesis would also imply that the β-GGM biosynthesis pathway has evolved to converge on a glucomannan structure closely related to XyG, an idea that raises interesting questions about the importance of this structure for function of both of the polysaccharides.

The molecular structure of hemicellulose polysaccharides influences their solubility and ability to interact with other cell wall components in ways that are not fully understood. It is notable that β-GGM has similarities in structure to XyG, suggesting their backbones and arrangements of branches confer beneficial properties. One distinguishing feature of β-GGM over the previously described AcGGM is the possession of disaccharide branches. What could be the advantage of this structure? The side chains might affect binding to cellulose in the cell wall. *In vitro* assays showed branches influence the XyG-bacterial cellulose interactions (Lopez et al., 2010), however, there is no clear evidence of an influence on XyG binding in plant cell walls. Second, the side chains may be important for recognition by cell wall modifying enzymes such as XTHs and mannanases (Pena et al., 2004; Schröder et al., 2006; Li et al., 2013; Ishida and Yokoyama, 2022). Thirdly, these side chains might influence solubility of the polymers. The *mur3-1 xlt2* double mutant (with mostly non-substituted XyG composed of XXXG units) can be partially or fully rescued by the addition of D-Gal, L-Ara*f*, or L-Ara*p* at the second or third Xyl*p* residue. This suggests that the disaccharide substitution frequency of XyG is an important parameter for XyG function, but perhaps not the identity or position of the substituted chains (Schultink et al., 2013; Zhu et al., 2018). Thus, a large decrease in XyG substitution in *mur3-3* causes a phenotype, while the smaller decrease in XyG β-Gal in the *xlt2* mutant has no effect. The loss of the side chains may promote inappropriate intracellular interactions of XyG or β-GGM, leading to the formation of membrane aggregates and Golgi secretion disruption (Madson et al., 2003; Zhao et al., 2019). This hypothesis is supported by the fact that the *mur3* phenotype is rescued by plant growth in increased temperature (Shirakawa et al., 1998; Kong et al., 2015). We showed that loss of β-galactosylation of β-GGM exacerbates the *mur3* XyG galactosylation mutant phenotypes, indicating a role for this β-Gal disaccharide side chain.

We were able to show using ssNMR that β-GGM is relatively immobile in the cell wall, consistent with binding of this hemicellulose to cellulose. In spruce wood, the AcGGM was found by ssNMR to have close proximity to the cellulose surface. It was further suggested that AcGGM binds to the cellulose surface in a two-fold screw conformation distinct from the soluble AcGGM conformation (Terrett et al., 2019). Here, based on the similarity of ^13^C NMR chemical shifts, we found no evidence for a change in conformation of the β-GGM between solution or in the intact cell wall. Recent molecular dynamics simulations of glucomannan suggest that the backbone Glc residues may promote maintenance of glycosidic bond angles consistent with a two-fold screw, through inter-residue H-bonding as seen in cellulose (Berglund et al., 2016; Berglund et al., 2019; Martinez-Abad et al., 2020). Perhaps a consequence of the disaccharide GM repeat is the maintenance of a flattened conformation, unlike that of the flexible conformation AcGGM which has relatively infrequent Glc residues. The simulations also suggested galactosylation of the Man residue further promotes the formation of two-fold screw ribbon conformation (Berglund et al., 2019). Thus, it is likely that the backbone of β-GGM in solution maintains a flattened conformation that can interact with cellulose without adopting a new shape.

We speculate that β-GGM is likely to interact with cellulose similarly to XyG, but with a few notable differences. XyG, with its glucan backbone, is able to interact with cellulose fibrils. Unlike xylan, which possesses a face that might dock into fibrils and hydrogen bond with the cellulose glucan chains (Busse-Wicher et al., 2016; Simmons et al., 2016; Grantham et al., 2017), XyG is thought to bind to the hydrophobic 100 or 200 cellulose fibril faces through stacking interactions and H-bonding, lying flat with substitutions placed on both sides of the two-fold screw backbone ribbon (Zhao et al., 2014; Benselfelt et al., 2016). Our earlier molecular dynamics simulations suggest β-GGM backbones, which contain alternating Man and Glc, could similarly bind to cellulose (Yu et al., 2018). Since the sugar backbone repeat is GM, in a two-fold screw ribbon each of the Man 2-OH that point out of the hexose ring plane could face away from the cellulose fibril. The β-GGM Glc residues would interact with cellulose as in the XyG backbone. The substitutions could additionally interact with the cellulose surface. However, since substitutions are only present on the alternating residues of Man in β-GGM, these will all lie on one side of the backbone ribbon, unlike XyG where substitutions will lie on both sides of the ribbon. This potentially provides somewhat different hemicellulose-cellulose interaction opportunities.

Studies of Arabidopsis seed mucilage give a hint that β-GGM does functionally interact with cellulose. Although the mucilage β-GGM differs in that it loses β-1,2-Gal at least in part through action of a cell wall β-galactosidase MUM2, the backbone and frequent α-Gal substitution of Man residues are typical of β-GGM. In mucilage, this β-GGM is important for arrangement of the cellulose, because the *csla2* and *magt1* mutants no longer form the normal cellulose rays as in WT (Yu et al., 2014; Voiniciuc et al., 2015). Indications of β-GGM influencing cellulose arrangements also comes from staining of the cellulose in etiolated hypocotyls, since altered arrangements were seen in the mutants lacking both β-GGM and XyG.

The structural similarity of β-GGM and XyG led us to hypothesize that they may play connected functions in the cell wall during growth and development. Previously, our knowledge of glucomannan function from Arabidopsis molecular genetics indicated a role limited to seed mucilage and in embryogenesis (Goubet et al., 2003; Goubet et al., 2009; Yu et al., 2014; Voiniciuc et al., 2015; Yu et al., 2018; Somssich et al., 2021). Our results support the idea that XyG conceals the importance of β-GGM in many tissues. For example, the Arabidopsis *csla2* β-GGM mutant shows few phenotypes in the plant, but it does have altered adherent mucilage (Yu et al., 2018). Notably, in the mucilage XyG is undetectable (Haughn and Western, 2012). Studies of the β-GGM and XyG backbone synthesis mutants also support a connection in function. The *csla2 xxt1 xxt2* mutant had more severe growth phenotypes than the *xxt1 xxt2* alone, again showing the role of β-GGM is partly obscured by XyG. We also showed that the loss of the β-GGM disaccharide side chain exacerbated the severity of XyG galactosylation mutant phenotypes, even though phenotypes were not observed in the presence of normal XyG. β-GGM and XyG are therefore connected in their functions, and they are both involved in cell expansion in various tissues. The relatively mild phenotypes of XyG mutants and β-GGM mutants is in part due to a level of functional redundancy of these hemicelluloses. It might be that loss of yet further hemicelluloses, including xylan, will reveal more severe impacts on wall function. The implication of our results is also that studies of XyG function have been hindered by the presence of β-GGM. β-GGM now needs to be studied alongside XyG in studies of hemicellulose function in plant cell expansion and development.

## Materials and Methods

### Plant Materials

Arabidopsis (*Arabidopsis thaliana*) plants used in this work were from Col-0 ecotype. The various mutants are: *mbgt1-1* (SALK_065561), *mbgt1-2* (SAIL_852_F05C), *csla2* (SALK_065083), *csla9* (SALK_071916), *magt1* (SALK_061576), *xxt1* (SAIL_785_E02), *xxt2* (SALK_101308), *mur3-1* (Reiter et al., *1997), mur3-3 (SALK_141953), xlt2* (GABI_552C10), *fut1* (*mur2-1*) (Reiter et al., 1997), *irx9l* (SALK_037323), *mum2-10* (SALK_011436), *csla2 clsa9* (Goubet et al., 2009), and *xxt1 xxt2* (Cavalier et al., 2008). The *csla2 xxt1 xxt2* triple mutant was generated by crossing *csla2* and *xxt1 xxt2*, the *irx9l xxt1 xxt2* triple mutant was generated by crossing *irx9l* and *xxt1 xxt2*, the *mbgt1-1 mur3-1* double mutant was generated by crossing *mbgt1-1* and *mur3-1*, and the *mbgt1-1 mur3-3* double mutant was generated by crossing *mbgt1-1* and *mur3-3*. The homozygous lines were identified by PCR. The primers used for genotyping are shown in Supplemental Table S3.

### Plant Growth Conditions

Plants were grown in controlled-environment chambers. Arabidopsis seeds were surface sterilized, sown on half Murashige and Skoog (MS) medium with 1% sucrose, stratified in darkness for 48 h at 4 °C, and then germinated at 21 °C under 16-h light/8-h dark conditions. After 10 days, the seedlings were transferred to soil and grown in growth chambers under the same conditions. Arabidopsis liquid callus cultures were generated and maintained as described in (Prime et al., 2000). Uniformly labelled ^13^C glucose was used in the medium to replace sucrose for ssNMR analysis.

*Nicotiana benthamiana* plants were grown at 21 °C under 16-h light/8-h dark conditions. Leaves of 4-week old *N. benthamiana* were used for infiltration.

Rosette leaves were harvested at 6 weeks, young stems at 30 days, siliques at 6 weeks, and mature stems at 8 weeks. The plant height, the number of rosette branches, and the number of cauline branches were measured at 7 and 8 weeks. A rosette branch was defined as one originating from axils on the unexpanded stem, while the cauline branch was defined as one originated from the expanded segment of the inflorescence stem (Keller et al., 2006). All experiments were performed on at least three independently harvested sets of plant material.

### Hypocotyl and Cell Measurements

Seeds were surface-sterilized, sown on MS plates, and stored at 4°C for 3 days. Seeds were exposed to light for 6 h to stimulate germination, then wrapped in two layers of aluminium foil and grown for 2 to 7 days at 21 °C. Plates with etiolated seedlings were scanned using an HP Scanjet 8300 scanner at 600 dpi, and hypocotyl length was measured using ImageJ. To measure cell length, 4-d-old etiolated seedlings were firstly analysed with Cryo-SEM. Four-day-old etiolated hypocotyls were mounted onto carbon pad stubs, frozen and then coated with platinum and maintained at −145 °C as described previously (Lyczakowski et al., 2019). Images were acquired on a Zeiss EVO HD15 using a backscattered electron detector and an accelerating voltage of 25 kV with a working distance of >15 mm. Cell length measurements were taken for cells at the base of the hypocotyl using ImageJ software.

For generating the heat maps comparing cell length between *xxt1 xxt2* and *csla2 xxt1 xxt2* mutants, four-day-old etiolated hypocotyls were submersed in 0.1 mg mL^-1^ propidium iodide for 3 minutes, washed briefly in water and then mounted in water on a microscope slide with a coverslip. The slide was mounted on an inverted Leica DMi8 SP8 confocal fitted with a 10x objective lens. Whole seedlings were imaged for fluorescence in 3D using the tile scan feature of the Leica LAS X Navigator software module and the tiles fused to generate a single z-stack file covering the whole hypocotyl region. The files were converted to tiff stacks and imported into MorphoGraphX (de Reuille et al., 2015). Voxels were averaged (XRad, YRad, ZRad = 2) and the following software tools implemented in this order: Edge detect (20,000), fill holes, closing, Marching cubes surface, located and deleted erroneous volumes manually, smooth mesh, subdivide, smooth mesh, project signal (5-10), Gaussian blur (2 px radius), draw seeds as long lines down the centre of each cell, watershed segmentation, corrected incorrect segmentations by drawing new seeds and resegmenting. An updated version of MorphoGraphX was obtained from Richard Smith (John Innes Centre, Norwich) which allows heat maps to be generated based on major axis length. These length heat maps were scaled from 0 to 200 micron range.

### Cellulose fluorescent staining and imaging

Four-day-old seedlings were stained according to the protocol described previously (Landrein et al., 2013). In our hands, cells in the upper portion of the hypocotyl stained uniformly while cells in the lower half did not. Expanded cells below the apical hook were therefore selected for imaging using an upright Leica SP8 confocal fitted with a 552 nm laser for excitation and 63x 1.4 NA oil immersion lens for imaging. Confocal optical sections were taken that covered the full depth of staining. Representative images in Figure 5 are taken from the middle of the upper cell wall surface with two consecutive sections averaged to aid observations of cellulose patterns. Orthogonal views were created by drawing line regions of interest along the length of the centre of cells using ImageJ and then using the reslice option.

### Preparation of soluble hemicelluloses

Dry and clean seeds were shaken in dH_2_O in a tube for 30 min at 30 Hz in a Retsch MM400 mill. The seed suspension was centrifuged at 1,000 rpm for 1 min. The supernatant was harvested and the seeds were washed twice with dH_2_O to get naked seeds. The mucilage supernatants were collected and used for mucilage analysis. Callus was harvested and washed with ddH_2_O to remove medium. Alcohol Insoluble Residue (AIR) from stems, leaves, seed mucilage, naked seeds, siliques, callus and etiolated hypocotyls was prepared as previously described (Goubet et al., 2009; Yu et al., 2018). Thirty milligrams of AIR was treated with 2.5 mL of 4 M NaOH at room temperature (RT) for 1 h and centrifuged at 4000 rpm for 15 min. In order to neutralize the NaOH, prior to enzymatic digestion, the supernatants were loaded onto a PD-10 desalting column (GE Life-Science) and eluted with 50 mM ammonium acetate (pH 6.0) according to the manufacture instruction. The eluent contained the majority of the de-acetylated hemicelluloses and was aliquoted into tubes for 25 mannan reactions digestions or 50 XyG reactions digestions.

### Enzyme Digestions

For mannan analysis, the hemicelluloses eluted from PD-10 were digested with an excess of *Cellvibrio japonicus* Man26A (*Cj*Man26A) mannanase (University of Newcastle) or *Aspergillus nidulans* GH5 (*An*GH5) mannanase (Novozymes) in 50 mM ammonium acetate (pH=6.0) at 37 °C overnight. Mannanases were de-activated after digestion with a heat treatment. Mannanase products were then digested overnight with *Aspergillus niger* GH35 β-galactosidase (Megazyme) or *Cellvibrio mixtus* GH27 α-galactosidase (Prozomix) in 50 mM ammonium acetate (pH 6.0) at 37 °C to remove the β-Gal or α-Gal side chains. For sequential digestion, enzymes used were: *Aspergillus niger* GH3 β-glucosidase (Novozymes), and *Cellvibrio mixtus* GH5 β-mannosidase (University of Newcastle). The digestion conditions were 50 mM ammonium acetate (pH 6.0) at 37 °C for 4 h with excess enzymes to complete digestion. After each reaction, samples were boiled at 100 °C for 10 min to denature the enzyme. Samples were then dried at 60 °C *in vacuo*.

For XyG analysis, the eluted hemicellulose fractions were digested with an excess of *Paenibacillus pabuli* XG5 (*Pp*XG5) xyloglucanase (Novozymes) in 50 mM ammonium acetate (pH 6.0) at 37 °C for 18 h.

For xylan analysis, the eluted hemicellulose fractions were digested with an excess of *Neocallimastix patriciarum* GH11 (*Np*GH11) xylanase (Megazyme) as previously described (Mortimer et al., 2010).

### Monosaccharide Analysis of AIR by HPAEC-PAD

Fifty micrograms of AIR was hydrolysed in 2 M TFA at 121 °C for 1 h, dried *in vacuo* and resuspended in H_2_O. Inositol was added as the internal standard. Chromatography of the samples was performed using a CarboPac PA20 column as previously described. A standard mixture, containing 25 μM of sugar standards (L-fucose, L-rhamnose, L-arabinose, D-galactose, D-glucose, D-xylose and D-mannose), was run before each batch of samples.

### Oligosaccharide Fingerprint Analysis by PACE

Samples and (Man)_1-6_ standards (Megazyme) were derivatized with 8-aminonapthalene-1,3,6-tresulphonic acid (ANTS; Invitrogen) as described previously (Goubet et al., 2002). After drying, the samples were re-suspended in 100 μl of 3 M urea, of which 2 μl was loaded onto the PACE gels. The samples were run and visualized using a G-box equipped with a trans-illuminator with long-wavelength light tubes (365 nm) and a short pass filter (500-600 nm) as described previously (Goubet et al., 2002). All analyses of oligosaccharides were repeated a minimum of three times.

### Preparation of Oligosaccharides for MS

Following the enzyme digestion, released peptides and enzymes were removed using reverse-phase Sep-Pak C18 cartridges (Waters) as previously described. The oligosaccharides were reductively aminated with 2-aminobenzamide (2-AB), using optimized labelling conditions. The labelled samples were then purified from reductive amination reagents using a GlycoClean S cartridge (Prozyme) as described previously (Tryfona et al., 2012).

### Hydrophilic Interaction Liquid Chromatography (HILIC)-MALDI-ToF MS/MS

Capillary HILIC was carried out using an LC-Packings Ultimate system (Dionex), using optimized elution conditions and robot harvest systems. After air drying, the sample spots were overlaid with 2,5-dihydroxybenzoic acid matrix and analysed by MALDI-ToF/ToF-MS/MS as described previously (Tryfona et al., 2012).

### Separation of oligosaccharides by SEC

Arabidopsis young stem AIR (500 mg), hydrolysed with an excess of enzymes (first *Cj*Man26A, and then by a combination of β-glucosidase, β-mannosidase, and α-galactosidase), was prepared as described above, and lyophilised. Samples were re-suspended in 2 ml dH_2_O, loaded onto a gravity-derived preparative Bio-Gel P2 column (190 × 2.5 cm; Bio-Rad), equilibrated and run in 20 mM ammonium acetate pH 6.0. Fractions were collected and dried *in vacuo*. Fraction of interest was determined by PACE.

### Solution-state NMR

Following SEC, lyophilised samples were re-suspended in D_2_O (700 μL; 99.9% purity) and transferred to a 5 mm NMR tube. NMR spectra were recorded at 298 K with a Bruker AVANCE III spectrometer operating at 600 MHz equipped with a TCI CryoProbe. ^1^H chemical-shift assignments were primarily obtained using ^1^H–^1^H total correlation spectroscopy (TOCSY) and rotating frame Overhauser effect spectroscopy (ROESY). The H-1/H-2 peaks in a double quantum filtered correlation spectroscopy (DQFCOSY) were used to remove ambiguities in the assignments of H-2. ^13^C assignments were obtained using ^13^C HSQC and H2BC experiments (although the latter was incomplete due to the low concentration of the sample) (Cavanagh et al., 1995; Nyberg et al., 2005); the mixing times were 70 and 200 msec for the TOCSY and ROESY experiments, respectively. Chemical shifts were measured relative to internal acetone (δ(^1^H) = 2.225, δ(^13^C) = 31.07 ppm). Data were processed using the Azara suite of programs and chemical-shift assignment was performed using CCPN Analysis v2.4 (Vranken et al., 2005).

### Protein expression and western blot analysis

3×Myc tagged *MBGT* (At4g13990) and *PgGUX* coding sequences were PCR amplified from synthetic DNA (IDT) or previously described constructs (Lyczakowski et al., 2017) using primers described in Supplemental Table S3. Tobacco infiltration, microsome isolation and western blot analysis of membrane preparations were all performed as previously described (Lyczakowski et al., 2017).

### β-Galactosyltransferase Activity Assay

Adherent mucilage hemicelluloses from WT seeds, rich in β-GGM lacking the β-gal, were prepared as previously described (Yu et al., 2018). *mbgt-1* soluble hemicellulose from young stem was prepared as above. WT adherent mucilage hemicelluloses and *mbgt-1* young stem hemicelluloses aliquots were dried and used as acceptors for *in vitro* β-Gal transfer reaction. UDP-Gal (5 mM) was replaced with water in certain reactions to control for non-specific galactosylation. Reaction was performed for 5 hours at room temperature and was terminated by heating the samples at 100 °C for 10 mins. The polysaccharides were extracted using methanol and chloroform as previously described (Lyczakowski et al., 2017). Extracted polysaccharides were digested with *Cj*Man26A and analysed with PACE.

### Phylogeny

The bulk of the GT47 Clade A sequences were downloaded as an orthologous cluster from the comparative genomics platform Plaza Dicots 4.5, Plaza Monocots 4.5 (Van Bel et al., 2018), and Plaza Gymnosperms 3.0 (Proost et al., 2015), but were supplemented with the results of HMMER (http://hmmer.org/) and TBLASTN (Altschul et al., 1990; Altschul et al., 1997) searches of additional published genomes (Hori et al., 2014; Filiault et al., 2018; Li et al., 2018; Weston et al., 2018; Chen et al., 2019; Zhang et al., 2020) using an Arabidopsis GT47 Clade A HMM or the *At*MUR3 protein sequence as a query, respectively. For the GT47-A tree, sequences were aligned with MAFFT (Katoh et al., 2002; Katoh and Standley, 2013) and truncated to their predicted GT47 domain (corresponding to residues 156–539 of *At*MUR3) using a custom Python script (https://www.python.org/). Substantially truncated and very poorly aligned sequences were removed from the alignment manually. Prottest3 (Darriba et al., 2011) was used to determine an appropriate substitution model (LG), and the tree was built with FastTreeMP (Price et al., 2010) with 100 bootstraps. For the smaller MBGT homologue tree, a subset of sequences was selected from the relevant subclade of the larger tree and aligned using MUSCLE (Edgar, 2004). Prottest3 indicated JTT+I+G+F to be the best substitution model, and the tree was built accordingly using RAxML (Stamatakis, 2014) with 100 rapid bootstraps; *At*MUR3 was included to root the tree as well as supplementary MBGT homologues from the genomes of *Nicotiana tabacum* (Sierro et al., 2014) and *Rosa chinensis* (Raymond et al., 2018).

### Preparation of callus sample for ssNMR

^13^C labelled callus was harvested and washed 6 times with unlabelled callus medium to remove the ^13^C glucose. Then the callus was frozen in liquid N_2_ and stored at −80 °C overnight. Frozen callus was ground into powder in liquid N_2_, thawed on ice and centrifuged at 15,000 rpm at 4 °C, removing excess liquid, twice to obtain moist callus sample for ssNMR.

### Solid-state NMR

Solid-state MAS NMR experiments were performed using Bruker (Karlsruhe, Germany) AVANCE NEO solid-state NMR spectrometers, operating at ^1^H and ^13^C Larmor frequencies of 1000.4 MHz and 251.6 MHz and 850.2 and 213.8 MHz, respectively, with 3.2 mm double-resonance E^free^ MAS probes. Experiments were conducted at an indicated temperature of 283 K at an MAS frequency of 12.5 kHz on both spectrometers. The ^13^C chemical shift was determined using the carbonyl peak at 177.8 ppm of L-alanine as an external reference with respect to tetramethylsilane (TMS). Both ^1^H−^13^C cross-polarisation (CP), with ramped (70–100%) ^1^H rf amplitude and 1 ms contact time, and direct polarisation (DP) were used to obtain the initial transverse magnetisation (Metz et al., 1994). While CP emphasises the more rigid material a short, 2 s, recycle delay DP experiment was used to preferentially detect the mobile components. Two-dimensional double-quantum (DQ) correlation spectra were recorded using the refocused INADEQUATE pulse sequence which relies upon the use of isotropic, scalar J coupling to obtain through-bond information regarding directly coupled nuclei (Lesage et al., 1997; Lesage et al., 1999; Fayon et al., 2005). The carbon 90° and 180° pulse lengths were 3.5 – 4.3 μs and 7.0 – 8.6 μs, respectively with 2τ spin-echo evolution times for a (π–τ–π/2) spin-echo of 4.48 ms. SPINAL-64 ^1^H decoupling was applied during both the evolution and signal acquisition periods at a ^1^H nutation frequency of 70–80 kHz (Fung et al., 2000). The acquisition time in the indirect dimension (*t*_1_) was 5.0 – 6.0 ms for the CP-INADEQUATE and 5.5 ms for the DP INADEQUATE experiment. The spectral width in the indirect dimension was 50 kHz for both with 192-416 acquisitions per *t*_1_ FID for the CP-INADEQUATE and 80 acquisitions for the DP INADEQUATE experiments. The States-TPPI method was used to achieve sign discrimination in *F*_1_. The recycle delay was 2 s for both CP INADEQUATE and DP INADEQUATE experiments. The spectra were obtained by Fourier transformation into 4 K (*F*_2_)× 2K (*F*_1_) points with exponential line broadening in *F*_2_ of 50 Hz for CP and 20 Hz for DP experiments respectively and squared sine bell processing in *F*_1_. All spectra obtained were processed and analysed using Bruker Topspin version 3.6.2.

### Accession Numbers

Sequence data from this article can be found in the Arabidopsis Genome Initiative or GenBank/EMBL databases under the following accession numbers: At4g13990 (MBGT1), At5g22740 (CSLA2), At5g03760 (CSLA9), At2g22900 (MAGT1), At3g62720 (XXT1), At4g02500 (XXT2), At1g27600 (IRX9L), At2g20370 (MUR3), At5g62220 (XLT2), At2g03220 (FUT1), and MUM2 (At5g63800).

## Author Contributions

Author contributions: L.Y., and P.D. conceived and designed the study. L.Y. conducted most of the experiments and analysed the data. X.Y. made callus for ssNMR. R.C., R.D. and S.P.B. conducted the solid-state NMR experiments. ssNMR data were analysed by R.C., R.D., Y.Y., L.Y., and P.D.. J.J.L. made the constructs and contributed to the MBGT protein expression for *in vitro* assay. R.W. performed the microscopy. L.F.L.W. did the phylogenetic analysis and statistics. Y.Y. measured the plant growth phenotypes. K.I. performed some of the PACE analysis. X.Y. and J.W-R. performed the crosses of plants. K.S. performed the solution NMR and analysed the data. K.B.R.M.K. assisted with guidance on the use of glycoside hydrolase enzymes and S.C. tested the enzyme specificities. O.M.T. and H.T. contributed to the data interpretation and project discussion. L.Y., Y.Y., and P.D. wrote the manuscript. J.J.L., L.F.L.W., Y.Y., O.M.T., and K. I. assisted writing the manuscript. All authors commented on and approved the final manuscript.

## Acknowledgements

We would like to acknowledge Prof. George Lomonossoff (John Innes Centre, UK), who developed the pEAQ-HyperTrans expression system used in this study. Plant Bioscience Limited supplied the pEAQ-HT vector that was used in this work. *An*GH5 mannanase, *An*GH3 β-glucosidase, and *Pp*XG5 xyloglucanase were kindly provided by Novozymes A/S, Denmark. *Cj*Man26A mannanase and *Cm*GH5 β-mannosidase were kind gifts from Harry Gilbert (University of Newcastle).

The Microscopy Core Facility at the Sainsbury Laboratory (Cambridge University) is supported by the Gatsby Charitable Foundation.

The UK High-Field Solid-State NMR Facility used in this research was funded by EPSRC and BBSRC (EP/T015063/1 and EP/R029946/1), as well as the University of Warwick including via part funding through Birmingham Science City Advanced Materials Projects 1 and 2 supported by Advantage West Midlands (AWM) and the European Regional Development Fund (ERDF). We wish to thank the Facility Manager Team (Dr Dinu Iuga and Dr Trent Franks, University of Warwick) for their help.

This work was supported by the Leverhulme Trust Centre for Natural Material Innovation and by Biotechnology and Biological Sciences Research Council (BBSRC) of the UK as part of the OpenPlant Synthetic Biology Research Centre (Reference BB/L014130/1), Cambridge BBSRC-DTP Programme (Reference BB/J014540/1), Broodbank Research Fellowship of University of Cambridge (no. PD16178) and a BBSRC iCASE studentship (Reference BB/M015432/1).

## Supplemental material Legends

**Supplemental Figure S1. Schematic structures of primary cell wall hemicellulose**.

**Supplemental Figure S2. PACE gels of control samples of un-digested material and enzymes**. A, un-digested control of etiolated seedling and naked seed samples. B, un-digested control of young stem samples. C, un-digested control of young stem samples of XyG related mutants. D, Background bands brought by enzymes used in this work, supporting that all PACE results were not contamination from enzymes themselves. Markers M to M_6_ are shown.

**Supplemental Figure S3. Structural analysis of α-galactosylated mannan oligosaccharides from *csla9* young stem**. A, Analysis of α-galactosylated mannan by PACE. *csla9* young stem was digested with *Cj*Man26A first and the resultant oligosaccharides were digested sequentially with α-galactosidase (α-Gal), β-glucosidase (β-Glc), and β-mannosidase (β-Man) enzymes. α-Galactosylated mannan oligosaccharides have Glc-Man repeating units. Markers M to M_6_ are shown. B, Hex5 (S1) in Figure 2B was analysed by high-energy CID MS/MS. The first Man at the non-reducing end is decorated with a single α-1,6-Gal.

**Supplemental Figure S4. Patterned β-GGM is widely present in eudicots**. A to C, Mannan from tomato fruit, kiwi fruit, and apple fruit was analysed. *An*GH5 products of AIR were digested with β-galactosidase (β-Gal) to test the presence of β-GGM. NC indicates a negative control without enzyme. D, Arabidopsis seed mucilage from WT and *mum2* was digested with *Cj*Man26A. β-GGM was detected in *mum2* mucilage. M, Man; G, Glc; Markers M to M_6_ are shown.

**Supplemental Figure S5. Loss of XyG does not affect the production of CSLA2 β-GGM, or *vice versa***. Five-week-old young stem was used for the following digestion. A, Assignment of *Pp*XG5 products of XyG digestion with PACE. The assignment is enabled by XyG-related mutants. B, Structure of XyG and β-GGM in *mur3-3* or *mbgt1-1* mutants. C, Structure of XyG and β-GGM in *xxt1 xxt2* and *csla2* mutants. Bands in red dashed lines are XyG oligosaccharide dimers. M: Man; G: Glc. Markers M to M_6_ are shown.

**Supplemental Figure S6. Arabidopsis callus hemicelluloses analyzed by PACE**. A, Callus mannan was analysed by both *Cj*Man26A and *An*GH5 mannanases. *csla2*-related mutants lack mannan accessed by *Cj*Man26A and *An*GH5. B, Callus XyG was analysed by *Pp*XG5. C, Callus xylan was analyzed by *Np*GH11. *irx9l*-related mutants lack xylan accessed by *Np*GH11. Bands in red dashed lines are XyG oligosaccharide dimers. M: Man; X: Xyl. Markers M to M_6_ and X to X_6_ are shown.

**Supplemental Figure S7. Solid-state NMR of WT Arabidopsis callus**. The carbohydrate region of a refocussed CP-INADEQUATE ^13^C MAS NMR spectrum of ^13^C enriched WT callus with the β-GGM Man peaks M5 and M6 labelled. Xyloglucan is also labelled as well as carbons in the major polysaccharides: galacturonic acid (GalA), terminal Xyl (X) and α-Gal and two arabinoses (A^t^ and A^c^). The terminal arabinose is labelled t and the other arabinose c. For cellulose, the environments have been split into two groups, domain 1 and 2 cellulose (C^1^ and C^2^). For xyloglucan, 5 sets of environments are seen depending on the substitution, labelled as unsubstituted backbone Glc (XyC^u^), substituted backbone Glc (XyC^s^), terminal Xyl on XyG (XgX^t^), substituted Xyl (XgX^s^), and Fuc (F). Assignments are listed in Supplementary Table 1. The inset shows an overlay for the M5, M6 region of a CP INADEQUATE spectrum of ^13^C enriched *irx9l xxt1 xxt2* callus (blue) with that of *csla2 xxt1 xxt2* callus. As expected, the β-GGM (M and α-Gal) peaks are missing from the *csla2 xxt1 xxt2* spectrum. Spectra were acquired at a ^13^C Larmor frequency of 213.8 MHz for WT callus and *csla2 xxt1 xxt2* and 251.6 MHz for *irx9l xxt1 xxt2*. The MAS frequency was 12.5 kHz and the spin-echo duration was 2.24 ms.

**Supplemental Table 1**. ^1^H and ^13^C NMR chemical shifts for the GGM oligosaccharide in solution and the solid-state NMR assignments of β-GGM.

**Supplemental Table 2**. Plant species used for the phylogenetic tree.

**Supplemental Table 3**. Primers used in this study.

## Parsed Citations

Altschul, S.F., Gish, W., Miller, W., Myers, E.W., and Lipman, D.J. (1990). Basic local alignment search tool. J Mol Biol 215, 403–410.

Altschul, S.F., Madden, T.L., Schaffer, A.A., Zhang, J., Zhang, Z., Miller, W., and Lipman, D.J. (1997). Gapped BLAST and PSI-BLAST: a new generation of protein database search programs. Nucleic Acids Res 25, 3389–3402.

Arnling Bååth, J., Martinez-Abad, A., Berglund, J., Larsbrink, J., Vilaplana, F., and Olsson, L. (2018). Mannanase hydrolysis of spruce galactoglucomannan focusing on the influence of acetylation on enzymatic mannan degradation. Biotechnol Biofuels 11, 114.

Aryal, B., Jonsson, K., Baral, A., Sancho-Andres, G., Routier-Kierzkowska, A.L., Kierzkowski, D., and Bhalerao, R.P. (2020). Interplay between Cell Wall and Auxin Mediates the Control of Differential Cell Elongation during Apical Hook Development. Curr Biol 30, 1733–1739.

Benselfelt, T., Cranston, E.D., Ondaral, S., Johansson, E., Brumer, H., Rutland, M.W., and Wågberg, L. (2016). Adsorption of xyloglucan onto cellulose surfaces of different morphologies: an entropy-driven process. Biomacromolecules 17, 2801–2811.

Berglund, J., Azhar, S., Lawoko, M., Lindström, M., Vilaplana, F., Wohlert, J., and Henriksson, G. (2019). The structure of galactoglucomannan impacts the degradation under alkaline conditions. Cellulose 26, 2155–2175.

Berglund, J., Angles d’Ortoli, T., Vilaplana, F., Widmalm, G., Bergenstråhle-Wohlert, M., Lawoko, M., Henriksson, G., Lindström, M., and Wohlert, J. (2016). A molecular dynamics study of the effect of glycosidic linkage type in the hemicellulose backbone on the molecular chain flexibility. Plant J 88, 56–70.

Bootten, T.J., Harris, P.J., Melton, L.D., and Newman, R.H. (2004). Solid-state 13C-NMR spectroscopy shows that the xyloglucans in the primary cell walls of mung bean (Vigna radiata L.) occur in different domains: a new model for xyloglucan-cellulose interactions in the cell wall. J Exp Bot 55, 571–583.

Burton, R.A., Gidley, M.J., and Fincher, G.B. (2010). Heterogeneity in the chemistry, structure and function of plant cell walls. Nat Chem Biol 6, 724–732.

Busse-Wicher, M., Grantham, N.J., Lyczakowski, J.J., Nikolovski, N., and Dupree, P. (2016). Xylan decoration patterns and the plant secondary cell wall molecular architecture. Biochem Soc T 44, 74–78.

Cavalier, D.M., Lerouxel, O., Neumetzler, L., Yamauchi, K., Reinecke, A., Freshour, G., Zabotina, O.A., Hahn, M.G., Burgert, I., Pauly, M., Raikhel, N.V., and Keegstra, K. (2008). Disrupting two Arabidopsis thaliana xylosyltransferase genes results in plants deficient in xyloglucan, a major primary cell wall component. Plant Cell 20, 1519–1537.

Cavanagh, J., Fairbrother, W.J., Palmer III, A.G., and Skelton, N.J. (1995). Protein NMR spectroscopy: principles and practice. (Elsevier).

Chen, J.H., Hao, Z.D., Guang, X.M., Zhao, C.X., Wang, P.K., Xue, L.J., Zhu, Q.H., Yang, L.F., Sheng, Y., Zhou, Y.W., Xu, H.B., Xie, H.Q., Long, X.F., Zhang, J., Wang, Z.R., Shi, M.M., Lu, Y., Liu, S.Q., Guan, L.H., Zhu, Q.H., Yang, L.M., Ge, S., Cheng, T.L., Laux, T., Gao, Q., Peng, Y., Liu, N., Yang, S.H., and Shi, J.S. (2019). Liriodendron genome sheds light on angiosperm phylogeny and species-pair differentiation. Nat Plants 5, 18–25.

Cocuron, J.C., Lerouxel, O., Drakakaki, G., Alonso, A.P., Liepman, A.H., Keegstra, K., Raikhel, N., and Wilkerson, C.G. (2007). A gene from the cellulose synthase-like C family encodes a β-1,4 glucan synthase. Proc Natl Acad Sci USA 104, 8550–8555.

Cosgrove, D.J. (2014). Re-constructing our models of cellulose and primary cell wall assembly. Curr Opin Plant Biol 22, 122–131.

Cosgrove, D.J. (2018). Nanoscale structure, mechanics and growth of epidermal cell walls. Curr Opin Plant Biol 46, 77–86.

Cresswell, R., Dupree, R., Brown, S.P., Pereira, C.S., Skaf, M.S., Sorieul, M., Dupree, P., and Hill, S. (2021). Importance of Water in Maintaining Softwood Secondary Cell Wall Nanostructure. Biomacromolecules 22, 4669–4680.

Darriba, D., Taboada, G.L., Doallo, R., and Posada, D. (2011). ProtTest 3: fast selection of best-fit models of protein evolution. Bioinformatics 27, 1164–1165.

de Reuille, P.B., Routier-Kierzkowska, A.-L., Kierzkowski, D., Bassel, G.W., Schüpbach, T., Tauriello, G., Bajpai, N., Strauss, S., Weber, A., and Kiss, A. (2015). MorphoGraphX: A platform for quantifying morphogenesis in 4D. Elife 4, e05864.

Dean, G.H., Zheng, H., Tewari, J., Huang, J., Young, D.S., Hwang, Y.T., Western, T.L., Carpita, N.C., McCann, M.C., Mansfield, S.D., and Haughn, G.W. (2007). The Arabidopsis MUM2 gene encodes a β-galactosidase required for the production of seed coat mucilage with correct hydration properties. Plant Cell 19, 4007–4021.

Edgar, R.C. (2004). MUSCLE: multiple sequence alignment with high accuracy and high throughput. Nucleic Acids Res 32, 1792–1797.

Fayon, F., Massiot, D., Levitt, M.H., Titman, J.J., Gregory, D.H., Duma, L., Emsley, L., and Brown, S.P. (2005). Through-space contributions to two-dimensional double-quantum J correlation NMR spectra of magic-angle-spinning solids. J Chem Phys 122, 194313.

Filiault, D.L., Ballerini, E.S., Mandáková, T., Aköz, G., Derieg, N.J., Schmutz, J., Jenkins, J., Grimwood, J., Shu, S.Q., Hayes, R.D., Hellsten, U., Barry, K., Yan, J.Y., Mihaltcheva, S., Karafiátová, M., Nizhynska, V., Kramer, E.M., Lysak, M.A., Hodges, S.A., and Nordborg, M. (2018). The Aquilegia genome provides insight into adaptive radiation and reveals an extraordinarily polymorphic chromosome with a unique history. Elife 7.

Fry, S.C., York, W.S., Albersheim, P., Darvill, A., Hayashi, T., Joseleau, J.P., Kato, Y., Lorences, E.P., Maclachlan, G.A., Mcneil, M., Mort, A.J., Reid, J.S.G., Seitz, H.U., Selvendran, R.R., Voragen, A.G.J., and White, A.R. (1993). An unambiguous nomenclature for xyloglucan-derived oligosaccharides. Physiol Plant 89, 1–3.

Fung, B., Khitrin, A., and Ermolaev, K. (2000). An improved broadband decoupling sequence for liquid crystals and solids. J Magn Reson 142, 97–101.

Geddes, D., and Wilkie, K. (1972). A galactoglucomannan from the stem tissues of the aquatic moss Fontinalis antipyretica. Carbohyd Res 23, 349–357.

Geshi, N., Harholt, J., Sakuragi, Y., Jensen, J.K., and Scheller, H.V. (2018). Glycosyltransferases of the GT 47 family. Annu Plant Rev, 265–283.

Gilbert, H.J. (2010). The biochemistry and structural biology of plant cell wall deconstruction. Plant Physiol 153, 444–455.

Goubet, F., Jackson, P., Deery, M.J., and Dupree, P. (2002). Polysaccharide analysis using carbohydrate gel electrophoresis: A method to study plant cell wall polysaccharides and polysaccharide hydrolases. Anal Biochem 300, 53–68.

Goubet, F., Misrahi, A., Park, S.K., Zhang, Z.N., Twell, D., and Dupree, P. (2003). AtCSLA7, a cellulose synthase-like putative glycosyltransferase, is important for pollen tube growth and embryogenesis in Arabidopsis. Plant Physiol 131, 547–557.

Goubet, F., Barton, C.J., Mortimer, J.C., Yu, X.L., Zhang, Z.N., Miles, G.P., Richens, J., Liepman, A.H., Seffen, K., and Dupree, P. (2009). Cell wall glucomannan in Arabidopsis is synthesised by CSLA glycosyltransferases, and influences the progression of embryogenesis. Plant J 60, 527–538.

Grantham, N.J., Wurman-Rodrich, J., Terrett, O.M., Lyczakowski, J.J., Stott, K., Iuga, D., Simmons, T.J., Durand-Tardif, M., Brown, S.P., Dupree, R., Busse-Wicher, M., and Dupree, P. (2017). An even pattern of xylan substitution is critical for interaction with cellulose in plant cell walls. Nat Plants 3, 859–865.

Han, M., Liu, Y., Zhang, F., Sun, D., and Jiang, J. (2020). Effect of galactose side-chain on the self-assembly of xyloglucan macromolecule. Carbohydr Polym 246, 116577.

Haughn, G.W., and Western, T.L. (2012). Arabidopsis Seed Coat Mucilage is a Specialized Cell Wall that Can be Used as a Model for Genetic Analysis of Plant Cell Wall Structure and Function. Front Plant Sci 3, 64.

Hori, K., Maruyama, F., Fujisawa, T., Togashi, T., Yamamoto, N., Seo, M., Sato, S., Yamada, T., Mori, H., Tajima, N., Moriyama, T., Ikeuchi, M., Watanabe, M., Wada, H., Kobayashi, K., Saito, M., Masuda, T., Sasaki-Sekimoto, Y., Mashiguchi, K., Awai, K., Shimojima, M., Masuda, S., Iwai, M., Nobusawa, T., Narise, T., Kondo, S., Saito, H., Sato, R., Murakawa, M., Ihara, Y., Oshima-Yamada, Y., Ohtaka, K., Satoh, M., Sonobe, K., Ishii, M., Ohtani, R., Kanamori-Sato, M., Honoki, R., Miyazaki, D., Mochizuki, H., Umetsu, J., Higashi, K., Shibata, D., Kamiya, Y., Sato, N., Nakamura, Y., Tabata, S., Ida, S., Kurokawa, K., and Ohta, H. (2014). Klebsormidium flaccidum genome reveals primary factors for plant terrestrial adaptation. Nat Commun 5, 3978.

Ishida, K., and Yokoyama, R. (2022). Reconsidering the function of the xyloglucan endotransglucosylase/hydrolase family. J Plant Res 135, 145–156.

Jensen, J.K., Schultink, A., Keegstra, K., Wilkerson, C.G., and Pauly, M. (2012). RNA-Seq analysis of developing nasturtium seeds (Tropaeolum majus): identification and characterization of an additional galactosyltransferase involved in xyloglucan biosynthesis. Mol Plant 5, 984–992.

Katoh, K., and Standley, D.M. (2013). MAFFT multiple sequence alignment software version 7: improvements in performance and usability. Mol Biol Evol 30, 772–780.

Katoh, K., Misawa, K., Kuma, K., and Miyata, T. (2002). MAFFT: a novel method for rapid multiple sequence alignment based on fast Fourier transform. Nucleic Acids Res 30, 3059–3066.

Keller, T., Abbott, J., Moritz, T., and Doerner, P. (2006). Arabidopsis REGULATOR OF AXILLARY MERISTEMS1 controls a leaf axil stem cell niche and modulates vegetative development. Plant Cell 18, 598–611.

Kim, S.J., Chandrasekar, B., Rea, A.C., Danhof, L., Zemelis-Durfee, S., Thrower, N., Shepard, Z.S., Pauly, M., Brandizzi, F., and Keegstra, K. (2020). The synthesis of xyloglucan, an abundant plant cell wall polysaccharide, requires CSLC function. Proc Natl Acad Sci USA 117, 20316–20324.

Kong, Y.Z., Pena, M.J., Renna, L., Avci, U., Pattathil, S., Tuomivaara, S.T., Li, X.M., Reiter, W.D., Brandizzi, F., Hahn, M.G., Darvill, A.G., York, W.S., and O’Neill, M.A. (2015). Galactose-depleted xyloglucan is dysfunctional and leads to dwarfism in Arabidopsis. Plant Physiol 167, 1296–U1294.

Landrein, B., Lathe, R., Bringmann, M., Vouillot, C., Ivakov, A., Boudaoud, A., Persson, S., and Hamant, O. (2013). Impaired cellulose synthase guidance leads to stem torsion and twists phyllotactic patterns in Arabidopsis. Curr Biol 23, 895–900.

Lesage, A., Bardet, M., and Emsley, L. (1999). Through-bond carbon-carbon connectivities in disordered solids by NMR. J Am Chem Soc 121, 10987–10993.

Lesage, A., Auger, C., Caldarelli, S., and Emsley, L. (1997). Determination of through-bond carbon-carbon connectivities in solid-state NMR using the INADEQUATE experiment. J Am Chem Soc 119, 7867–7868.

Li, F.W., Brouwer, P., Carretero-Paulet, L., Cheng, S., de Vries, J., Delaux, P.M., Eily, A., Koppers, N., Kuo, L.Y., Li, Z., Simenc, M., Small, I., Wafula, E., Angarita, S., Barker, M.S., Brautigam, A., dePamphilis, C., Gould, S., Hosmani, P.S., Huang, Y.M., Huettel, B., Kato, Y., Liu, X., Maere, S., McDowell, R., Mueller, L.A., Nierop, K.G.J., Rensing, S.A., Robison, T., Rothfels, C.J., Sigel, E.M., Song, Y., Timilsena, P.R., Van de Peer, Y., Wang, H., Wilhelmsson, P.K.I., Wolf, P.G., Xu, X., Der, J.P., Schluepmann, H., Wong, G.K., and Pryer, K.M. (2018). Fern genomes elucidate land plant evolution and cyanobacterial symbioses. Nat Plants 4, 460–472.

Li, W.B., Guan, Q.M., Wang, Z.Y., Wang, Y.D., and Zhu, J.H. (2013). A bi-functional xyloglucan galactosyltransferase is an indispensable salt stress tolerance determinant in Arabidopsis. Mol Plant 6, 1344–1354.

Li, X.M., Cordero, I., Caplan, J., Molhoj, M., and Reiter, W.D. (2004). Molecular analysis of 10 coding regions from arabidopsis that are homologous to the MUR3 xyloglucan galactosyltransferase. Plant Physiol 134, 940–950.

Liepman, A.H., Wilkerson, C.G., and Keegstra, K. (2005). Expression of cellulose synthase-like (Csl) genes in insect cells reveals that CslAfamily members encode mannan synthases. Proc Natl Acad Sci USA 102, 2221–2226.

Liepman, A.H., Nairn, C.J., Willats, W.G., Sorensen, I., Roberts, A.W., and Keegstra, K. (2007). Functional genomic analysis supports conservation of function among cellulose synthase-like a gene family members and suggests diverse roles of mannans in plants. Plant Physiol 143, 1881–1893.

Liu, L., Paulitz, J., and Pauly, M. (2015). The presence of fucogalactoxyloglucan and its synthesis in rice indicates conserved functional importance in plants. Plant Physiol 168, 549–560.

Lopez, M., Bizot, H., Chambat, G., Marais, M.F., Zykwinska, A., Ralet, M.C., Driguez, H., and Buleon, A. (2010). Enthalpic studies of xyloglucan-cellulose interactions. Biomacromolecules 11, 1417–1428.

Lyczakowski, J.J., Bourdon, M., Terrett, O.M., Helariutta, Y., Wightman, R., and Dupree, P. (2019). Structural imaging of native cryo-preserved secondary cell walls reveals the presence of macrofibrils and their formation requires normal cellulose, lignin and xylan biosynthesis. Front Plant Sci 10, 1398.

Lyczakowski, J.J., Wicher, K.B., Terrett, O.M., Faria-Blanc, N., Yu, X., Brown, D., Krogh, K., Dupree, P., and Busse-Wicher, M. (2017). Removal of glucuronic acid from xylan is a strategy to improve the conversion of plant biomass to sugars for bioenergy. Biotechnol Biofuels 10, 224.

Macquet, A., Ralet, M.C., Loudet, O., Kronenberger, J., Mouille, G., Marion-Poll, A., and North, H.M. (2007). A naturally occurring mutation in an Arabidopsis accession affects a beta-D-galactosidase that increases the hydrophilic potential of rhamnogalacturonan I in seed mucilage. Plant Cell 19, 3990–4006.

Madson, M., Dunand, C., Li, X.M., Verma, R., Vanzin, G.F., Calplan, J., Shoue, D.A., Carpita, N.C., and Reiter, W.D. (2003). The MUR3 gene of Arabidopsis encodes a xyloglucan galactosyltransferase that is evolutionarily related to animal exostosins. Plant Cell 15, 1662–1670.

Martinez-Abad, A., Jimenez-Quero, A., Wohlert, J., and Vilaplana, F. (2020). Influence of the molecular motifs of mannan and xylan populations on their recalcitrance and organization in spruce softwoods. Green Chem 22, 3956–3970.

Metz, G., Wu, X.L., and Smith, S.O. (1994). Ramped-amplitude cross-polarization in magic-angle-spinning NMR. J Magn Reson Ser A 110, 219–227.

Mikkelsen, M.D., Harholt, J., Ulvskov, P., Johansen, I.E., Fangel, J.U., Doblin, M.S., Bacic, A., and Willats, W.G. (2014). Evidence for land plant cell wall biosynthetic mechanisms in charophyte green algae. Ann Bot 114, 1217–1236.

Mikkelsen, M.D., Harholt, J., Westereng, B., Domozych, D., Fry, S.C., Johansen, I.E., Fangel, J.U., Lezyk, M., Feng, T., Nancke, L., Mikkelsen, J.D., Willats, W.G.T., and Ulvskov, P. (2021). Ancient origin of fucosylated xyloglucan in charophycean green algae. Commun Biol 4, 754.

Mortimer, J.C., Miles, G.P., Brown, D.M., Zhang, Z., Segura, M.P., Weimar, T., Yu, X., Seffen, K.A., Stephens, E., Turner, S.R., and Dupree, P. (2010). Absence of branches from xylan in Arabidopsis gux mutants reveals potential for simplification of lignocellulosic biomass. Proc Natl Acad Sci U S A 107, 17409–17414.

Nikolovski, N., Rubtsov, D., Segura, M.P., Miles, G.P., Stevens, T.J., Dunkley, T.P.J., Munro, S., Lilley, K.S., and Dupree, P. (2012). Putative Glycosyltransferases and Other Plant Golgi Apparatus Proteins Are Revealed by LOPIT Proteomics. Plant Physiol 160, 1037–1051.

Nothnagel, A.L., and Nothnagel, E.A. (2007). Primary cell wall structure in the evolution of land plants. J Integr Plant Biol 49, 1271–1278.

Nyberg, N.T., Duus, J.O., and Sorensen, O.W. (2005). Heteronuclear two-bond correlation: suppressing heteronuclear three-bond or higher NMR correlations while enhancing two-bond correlations even for vanishing 2JCH. J Am Chem Soc 127, 6154–6155.

Obayashi, T., Aoki, Y., Tadaka, S., Kagaya, Y., and Kinoshita, K. (2018). ATTED-II in 2018: A Plant Coexpression Database Based on Investigation of the Statistical Property of the Mutual Rank Index. Plant Cell Physiol 59, 440.

Park, Y.B., and Cosgrove, D.J. (2015). Xyloglucan and its interactions with other components of the growing cell wall. Plant Cell Physiol 56, 180–194.

Pauly, M., and Keegstra, K. (2016). Biosynthesis of the plant cell wall matrix polysaccharide xyloglucan. Annu Rev Plant Biol 67, 235–259.

Pena, M.J., Kong, Y.Z., York, W.S., and O’Neill, M.A. (2012). A galacturonic acid-containing xyloglucan is involved in Arabidopsis root hair tip growth. Plant Cell 24, 4511–4524.

Pena, M.J., Ryden, P., Madson, M., Smith, A.C., and Carpita, N.C. (2004). The galactose residues of xyloglucan are essential to maintain mechanical strength of the primary cell walls in Arabidopsis during growth. Plant Physiol 134, 443–451.

Popper, Z.A., and Fry, S.C. (2003). Primary cell wall composition of bryophytes and charophytes. Ann Bot 91, 1–12.

Price, M.N., Dehal, P.S., and Arkin, A.P. (2010). FastTree 2–approximately maximum-likelihood trees for large alignments. Plos One 5.

Prime, T.A., Sherrier, D.J., Mahon, P., Packman, L.C., and Dupree, P. (2000). A proteomic analysis of organelles from Arabidopsis thaliana. Electrophoresis 21, 3488–3499.

Proost, S., Van Bel, M., Vaneechoutte, D., Van de Peer, Y., Inze, D., Mueller-Roeber, B., and Vandepoele, K. (2015). PLAZA 3.0: an access point for plant comparative genomics. Nucleic Acids Res 43, D974–981.

Raymond, O., Gouzy, J., Just, J., Badouin, H., Verdenaud, M., Lemainque, A., Vergne, P., Moja, S., Choisne, N., Pont, C., Carrère, S., Caissard, J.C., Couloux, A., Cottret, L., Aury, J.M., Szécsi, J., Latrasse, D., Madoui, M.A., Francois, L., Fu, X., Yang, S.H., Dubois, A., Piola, F., Larrieu, A., Perez, M., Labadie, K., Perrier, L., Govetto, B., Labrousse, Y., Villand, P., Bardoux, C., Boltz, V., Lopez-Roques, C., Heitzler, P., Vernoux, T., Vandenbussche, M., Quesneville, H., Boualem, A., Bendahmane, A., Liu, C., Le Bris, M., Salse, J., Baudino, S., Benhamed, M., Wincker, P., and Bendahmane, M. (2018). The Rosa genome provides new insights into the domestication of modern roses. Nat Genet 50, 772–777.

Reiter, W.D., Chapple, C., and Somerville, C.R. (1997). Mutants of Arabidopsis thaliana with altered cell wall polysaccharide composition. Plant J 12, 335–345.

Robert, M., Waldhauer, J., Stritt, F., Yang, B., Pauly, M., and Voiniciuc, C. (2021). Modular biosynthesis of plant hemicellulose and its impact on yeast cells. Biotechnol Biofuels 14, 140.

Rodrı’guez-Gacio, C., Iglesias-Ferna’ndez, R., Carbonero, P., and Matilla, A.J. (2012). Softening-up mannan-rich cell walls. J Exp Bot 63, 3976–3988.

Scheller, H.V., and Ulvskov, P. (2010). Hemicelluloses. Annu Rev Plant Biol 61, 263–289.

Schröder, R., Wegrzyn, T.F., Sharma, N.N., and Atkinson, R.G. (2006). LeMAN4 endo-β-mannanase from ripe tomato fruit can act as a mannan transglycosylase or hydrolase. Planta 224, 1091–1102.

Schröder, R., Nicolas, P., Vincent, S.J., Fischer, M., Reymond, S., and Redgwell, R.J. (2001). Purification and characterisation of a galactoglucomannan from kiwifruit (Actinidia deliciosa). Carbohydr Res 331, 291–306.

Schultink, A., Cheng, K., Park, Y.B., Cosgrove, D.J., and Pauly, M. (2013). The identification of two arabinosyltransferases from tomato reveals functional equivalency of xyloglucan side chain substituents. Plant Physiol 163, 86–94.

Shirakawa, M., Yamatoya, K., and Nishinari, K. (1998). Tailoring of xyloglucan properties using an enzyme. Food Hydrocoll 12, 25–28.

Sierro, N., Battey, J.N., Ouadi, S., Bakaher, N., Bovet, L., Willig, A., Goepfert, S., Peitsch, M.C., and Ivanov, N.V. (2014). The tobacco genome sequence and its comparison with those of tomato and potato. Nat Commun 5, 1–9.

Simmons, T.J., Mortimer, J.C., Bernardinelli, O.D., Pöppler, A.C., Brown, S.P., Deazevedo, E.R., Dupree, R., and Dupree, P. (2016). Folding of xylan onto cellulose fibrils in plant cell walls revealed by solid-state NMR. Nat Commun 7, 13902.

Sims, I.M., Craik, D.J., and Bacic, A. (1997). Structural characterisation of galactoglucomannan secreted by suspension-cultured cells of Nicotiana plumbaginifolia. Carbohydr Res 303, 79–92.

Somssich, M., Vandenbussche, F., Ivakov, A., Funke, N., Ruprecht, C., Vissenberg, K., Vander Straeten, D., Persson, S., and Suslov, D. (2021). Brassinosteroids influence arabidopsis hypocotyl graviresponses through changes in mannans and cellulose. Plant Cell Physiol 62, 678–692.

Stamatakis, A. (2014). RAxML version 8: a tool for phylogenetic analysis and post-analysis of large phylogenies. Bioinformatics 30, 1312–1313.

Tamura, K., Shimada, T., Kondo, M., Nishimura, M., and Hara-Nishimura, I. (2005). KATAMARI1/MURUS3 is a novel Golgi membrane protein that is required for endomembrane organization in Arabidopsis. Plant Cell 17, 1764–1776.

Tedman-Jones, J.D., Lei, R., Jay, F., Fabro, G., Li, X., Reiter, W.D., Brearley, C., and Jones, J.D. (2008). Characterization of Arabidopsis mur3 mutations that result in constitutive activation of defence in petioles, but not leaves. Plant J 56, 691–703.

Terrett, O.M., Lyczakowski, J.J., Yu, L., Iuga, D., Franks, W.T., Brown, S.P., Dupree, R., and Dupree, P. (2019). Molecular architecture of softwood revealed by solid-state NMR. Nat Commun 10, 4978.

Thomas, J., Idris, N.A., and Collings, D.A. (2017). Pontamine fast scarlet 4B bifluorescence and measurements of cellulose microfibril angles. J Microsc 268, 13–27.

Tryfona, T., Liang, H.C., Kotake, T., Tsumuraya, Y., Stephens, E., and Dupree, P. (2012). Structural characterization of arabidopsis leaf arabinogalactan polysaccharides. Plant Physiol 160, 653–666.

Van Bel, M., Diels, T., Vancaester, E., Kreft, L., Botzki, A., Van de Peer, Y., Coppens, F., and Vandepoele, K. (2018). PLAZA 4.0: an integrative resource for functional, evolutionary and comparative plant genomics. Nucleic Acids Res 46, D1190–D1196.

Velasquez, S.M., Guo, X., Gallemi, M., Aryal, B., Venhuizen, P., Barbez, E., Dunser, K.A., Darino, M., Pĕnčík, A., Novák, O., Kalyna, M., Mouille, G., Benkova, E., R, P.B., Mravec, J., and Kleine-Vehn, J. (2021). Xyloglucan remodeling defines auxin-dependent differential tissue expansion in plants. Int J Mol Sci 22.

Voiniciuc, C., Dama, M., Gawenda, N., Stritt, F., and Pauly, M. (2019). Mechanistic insights from plant heteromannan synthesis in yeast. P Natl Acad Sci USA 116, 522–527.

Voiniciuc, C., Schmidt, M.H., Berger, A., Yang, B., Ebert, B., Scheller, H.V., North, H.M., Usadel, B., and Günl, M. (2015). MUCILAGE-RELATED10 produces galactoglucomannan that maintains pectin and cellulose architecture in arabidopsis seed mucilage. Plant Physiol 169, 403–420.

von Freiesleben, P., Spodsberg, N., Blicher, T.H., Anderson, L., Jørgensen, H., Stålbrand, H., Meyer, A.S., and Krogh, K.B. (2016). An Aspergillus nidulans GH26 endo-β-mannanase with a novel degradation pattern on highly substituted galactomannans. Enzyme and microb technol 83, 68–77.

Vranken, W.F., Boucher, W., Stevens, T.J., Fogh, R.H., Pajon, A., Llinas, M., Ulrich, E.L., Markley, J.L., Ionides, J., and Laue, E.D. (2005). The CCPN data model for NMR spectroscopy: development of a software pipeline. Proteins: struct funct bioinform 59, 687–696.

Wang, S., Li, L., Li, H., Sahu, S.K., Wang, H., Xu, Y., Xian, W., Song, B., Liang, H., Cheng, S., Chang, Y., Song, Y., Cebi, Z., Wittek, S., Reder, T., Peterson, M., Yang, H., Wang, J., Melkonian, B., Van de Peer, Y., Xu, X., Wong, G.K., Melkonian, M., Liu, H., and Liu, X. (2020). Genomes of early-diverging streptophyte algae shed light on plant terrestrialization. Nat Plants 6, 95–106.

Wei, Q.Q., Yang, Y., Li, H., Liu, Z.W., Fu, R., Feng, H.Q., and Li, C. (2021). The xyloglucan galactosylation modulates the cell wall stability of pollen tube. Planta 254.

Weston, D.J., Turetsky, M.R., Johnson, M.G., Granath, G., Lindo, Z., Belyea, L.R., Rice, S.K., Hanson, D.T., Engelhardt, K.A., and Schmutz, J. (2018). The sphagnome project: enabling ecological and evolutionary insights through a genus-level sequencing project. New Phytol 217, 16–25.

Whitney, S.E.C., Wilson, E., Webster, J., Bacic, A., Reid, J.S.G., and Gidley, M.J. (2006). Effects of structural variation in xyloglucan polymers on interactions with bacterial cellulose. Am J Bot 93, 1402–1414.

Xiao, C., Zhang, T., Zheng, Y., Cosgrove, D.J., and Anderson, C.T. (2016). Xyloglucan deficiency disrupts microtubule stability and cellulose biosynthesis in arabidopsis, altering cell growth and morphogenesis. Plant Physiol 170, 234–249.

Yin, Y., Huang, J., and Xu, Y. (2009). The cellulose synthase superfamily in fully sequenced plants and algae. BMC Plant Biol 9, 99.

Yu, L., Lyczakowski, J.J., Pereira, C.S., Kotake, T., Yu, X.L., Li, A., Mogelsvang, S., Skaf, M.S., and Dupree, P. (2018). The patterned structure of galactoglucomannan suggests it may bind to cellulose in seed mucilage. Plant Physiol 178, 1011–1026.

Yu, L., Shi, D., Li, J., Kong, Y., Yu, Y., Chai, G., Hu, R., Wang, J., Hahn, M.G., and Zhou, G. (2014). CELLULOSE SYNTHASE-LIKE A2, a glucomannan synthase, is involved in maintaining adherent mucilage structure in Arabidopsis seed. Plant Physiol 164, 1842–1856.

Zhang, L.S., Chen, F., Zhang, X.T., Li, Z., Zhao, Y.Y., Lohaus, R., Chang, X.J., Dong, W., Ho, S.Y.W., Liu, X., Song, A.X., Chen, J.H., Guo, W.L., Wang, Z.J., Zhuang, Y.Y., Wang, H.F., Chen, X.Q., Hu, J., Liu, Y.H., Qin, Y., Wang, K., Dong, S.S., Liu, Y., Zhang, S.Z., Yu, X.X., Wu, Q., Wang, L.S., Yan, X.Q., Jiao, Y.N., Kong, H.Z., Zhou, X.F., Yu, C.W., Chen, Y.C., Li, F., Wang, J.H., Chen, W., Chen, X.L., Jia, Q.D., Zhang, C., Jiang, Y.F., Zhang, W.B., Liu, G.H., Fu, J.Y., Chen, F., Ma, H., Van de Peer, Y., and Tang, H.B. (2020). The water lily genome and the early evolution of flowering plants. Nature 577, 79–84.

Zhang, X.Y., Rogowski, A., Zhao, L., Hahn, M.G., Avci, U., Knox, J.P., and Gilbert, H.J. (2014). Understanding how the complex molecular architecture of mannan-degrading hydrolases contributes to plant cell wall degradation. J Biol Chem 289, 2002–2012.

Zhang, Y., Yu, J.Y., Wang, X., Durachko, D.M., Zhang, S.L., and Cosgrove, D.J. (2021). Molecular insights into the complex mechanics of plant epidermal cell walls. Science 372, 706–711.

Zhao, F., Chen, W., Sechet, J., Martin, M., Bovio, S., Lionnet, C., Long, Y., Battu, V., Mouille, G., Moneger, F., and Traas, J. (2019). Xyloglucans and microtubules synergistically maintain meristem geometry and phyllotaxis. Plant Physiol 181, 1191–1206.

Zhao, Z., Crespi, V.H., Kubicki, J.D., Cosgrove, D.J., and Zhong, L. (2014). Molecular dynamics simulation study of xyloglucan adsorption on cellulose surfaces: effects of surface hydrophobicity and side-chain variation. Cellulose 21, 1025–1039.

Zhu, L., Dama, M., and Pauly, M. (2018). Identification of an arabinopyranosyltransferase from Physcomitrella patens involved in the synthesis of the hemicellulose xyloglucan. Plant Direct 2, e00046.

