## Supplemental material for "Eudicot primary cell wall glucomannan is related in synthesis, structure and function to xyloglucan"

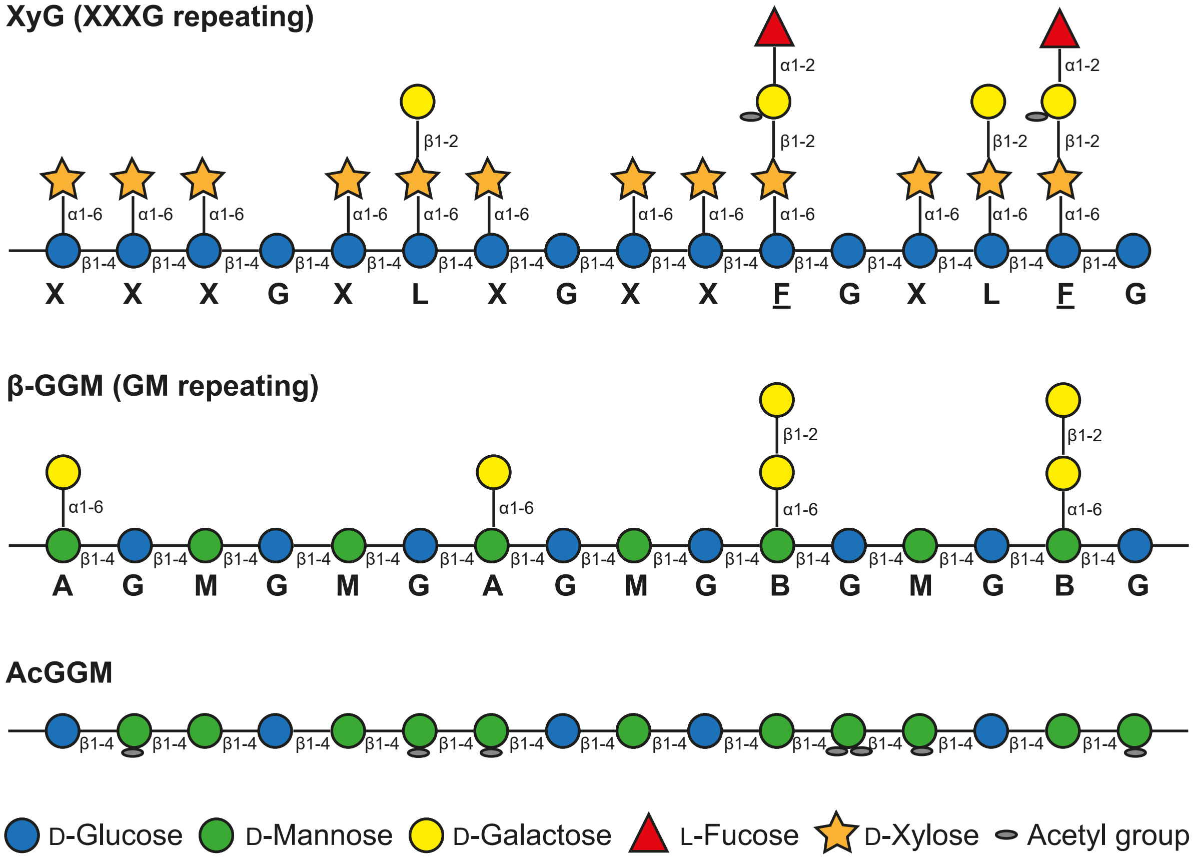

**Supplemental Figure S1. Schematic structures of primary cell wall hemicellulose.**

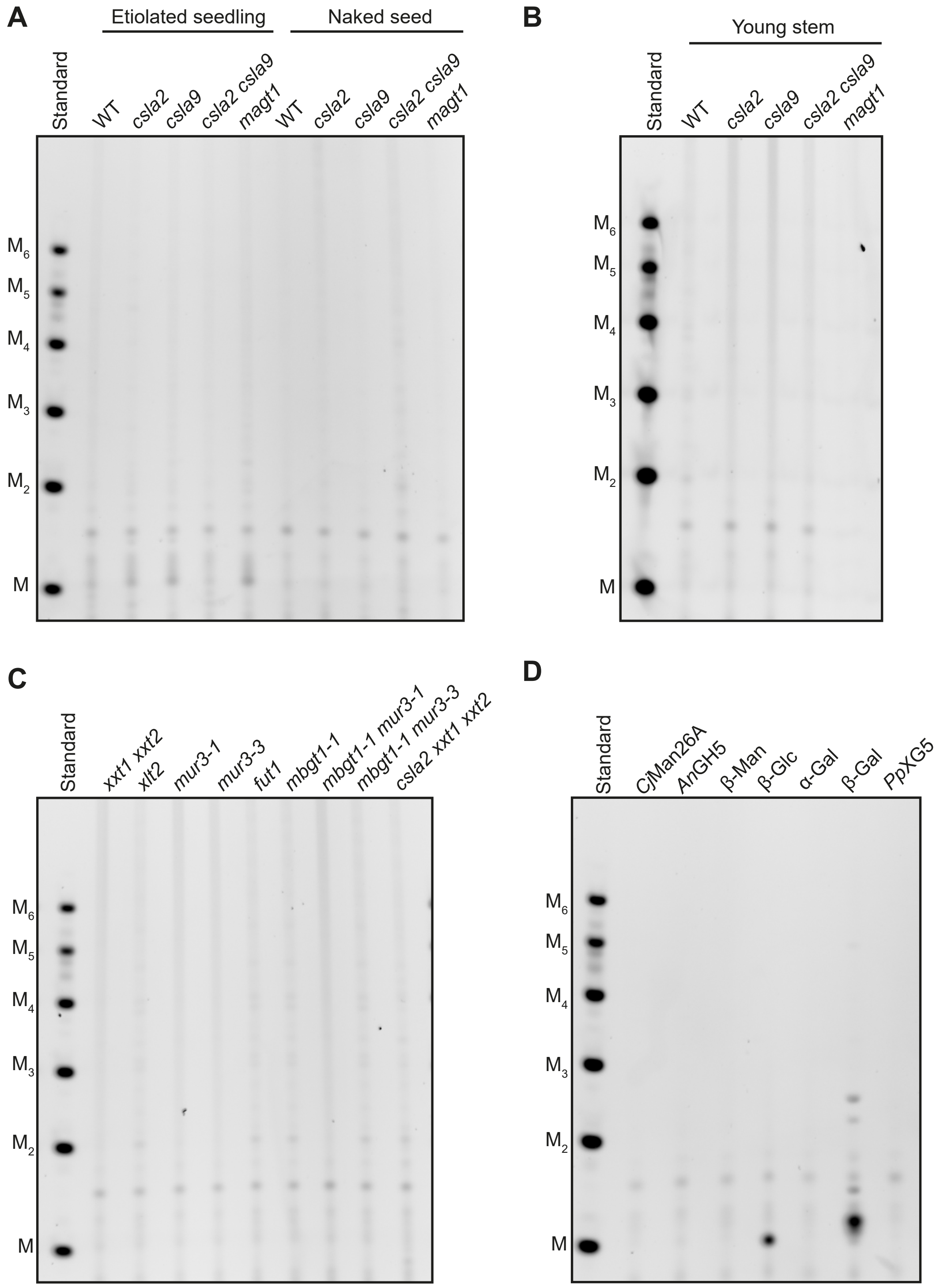

**Supplemental Figure S2. PACE gels of control samples of un-digested material and enzymes.** A, Un-digested control of etiolated seedling and naked seed samples. B, Un-digested control of young stem samples. C, Un-digested control of young stem samples of XyG related mutants. D, Background bands brought by enzymes used in this work, supporting that all PACE results were not contamination by enzymes themselves. Markers M to M6 are shown. Markers M to $M_6$ are shown.

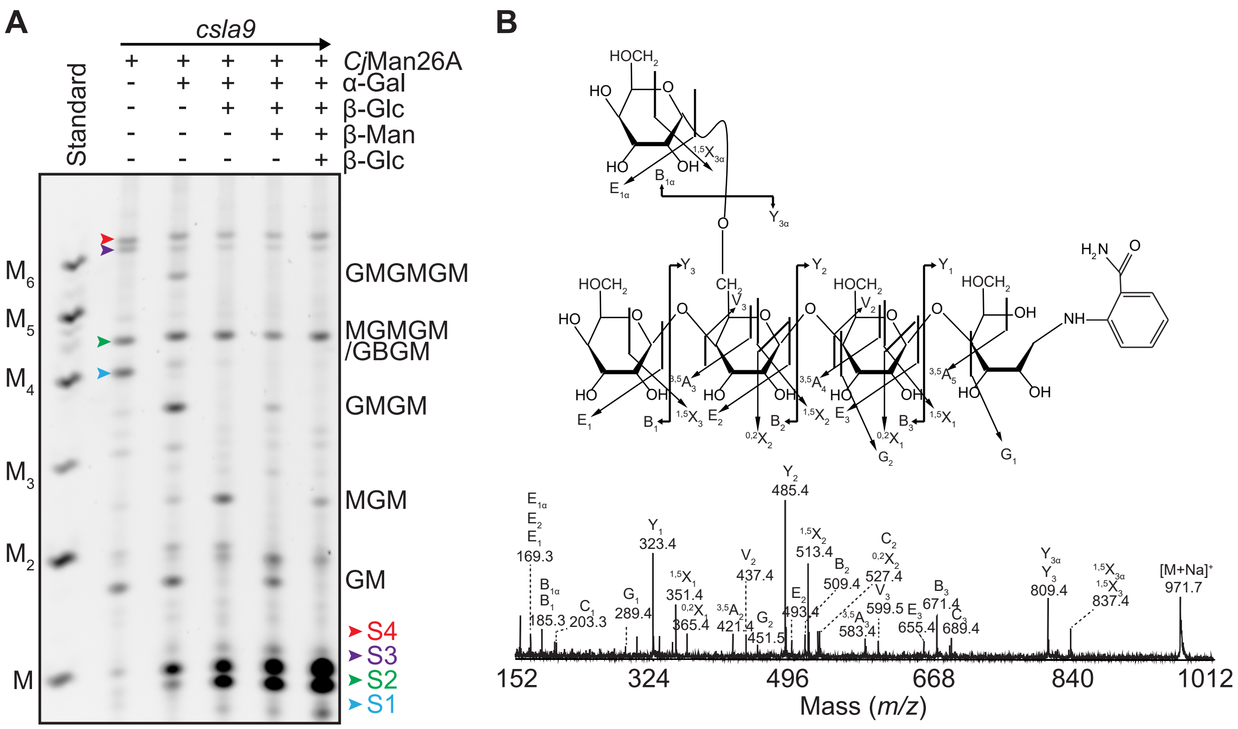

**Supplemental Figure S3. Structural analysis of α-galactosylated mannan oligosaccharides from *csla9* young stem.** A, Analysis of α-galactosylated mannan by PACE. *csla9* young stem was digested with *Cj*Man26A first and the resultant oligosaccharides were digested sequentially with α-galactosidase (α-Gal), β-glucosidase (β-Glc), and β-mannosidase (β-Man) enzymes. α-Galactosylated mannan oligosaccharides have Glc-Man repeating units. Markers M to $M_6$ are shown. B, Hex5 (S1) in Figure 2B was analysed by high-energy CID MS/MS. The first Man at the non-reducing end is decorated with a single α-1,6-Gal.

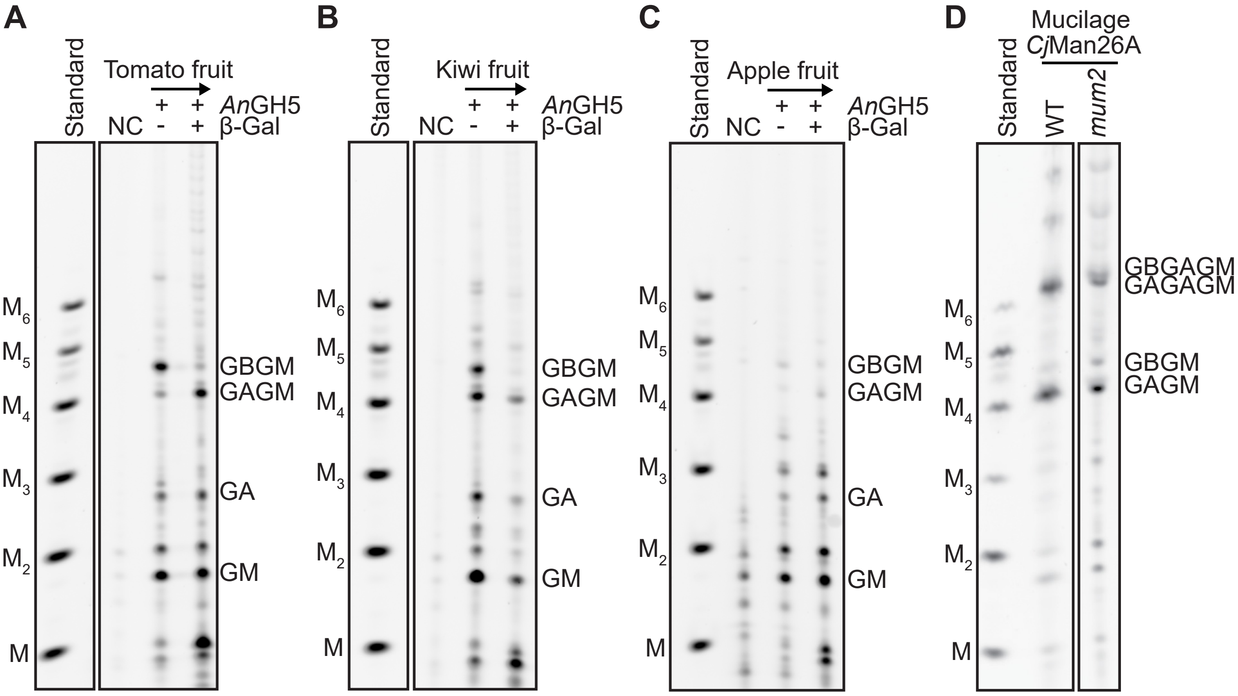

**Supplemental Figure S4. Patterned β-GGM is widely present in eudicots.** A to C, Mannans from tomato fruit, kiwi fruit, and apple fruit were analysed. *An*GH5 products of AIR were digested with β-galactosidase (β-Gal) to test the presence of β-GGM. NC indicates a negative control without enzyme. D, Arabidopsis seed mucilage from WT and *mum2* was digested with *Cj*Man26A. β-GGM was detected in *mum2* mucilage. M, Man; G, Glc; Markers M to $M_6$ are shown.

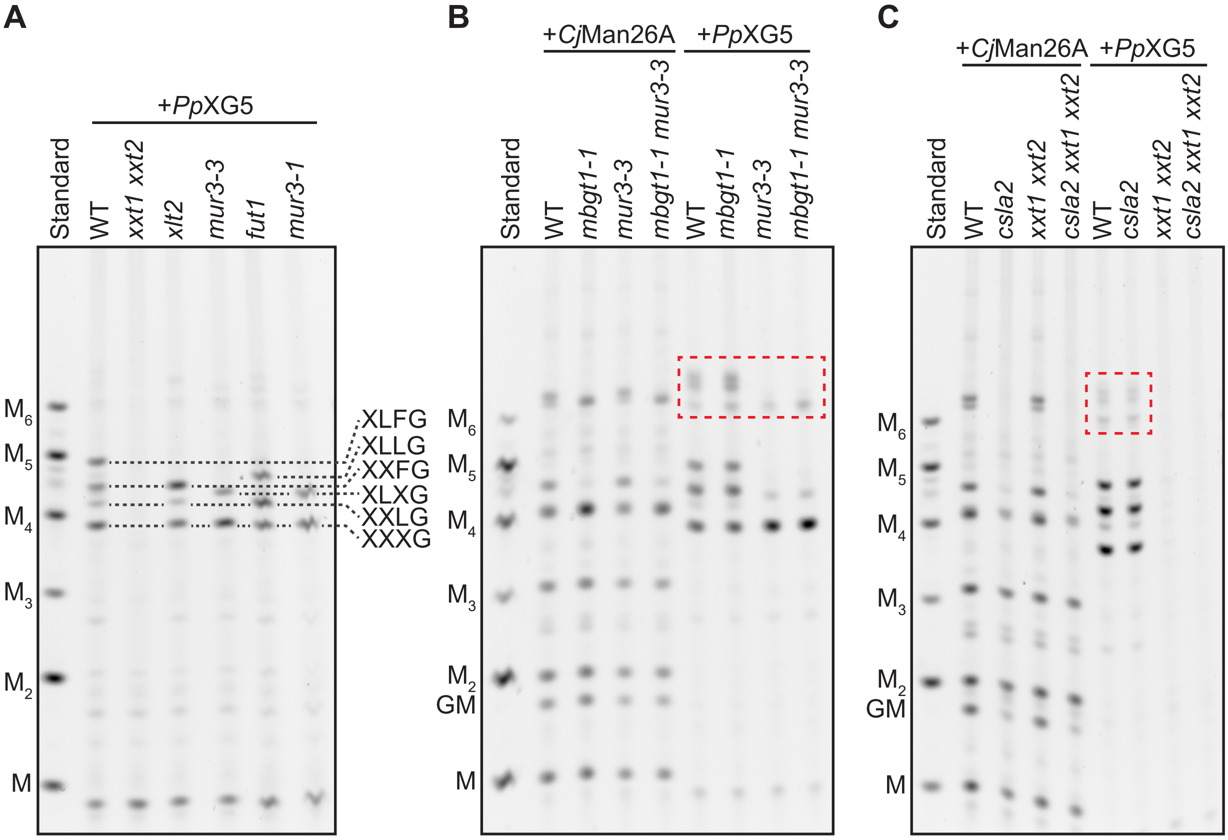

**Supplemental Figure S5. Loss of XyG does not affect the production of CSLA2 β-GGM, or vice versa.** Five-week-old young stem was used for the following digestion. A, Assignment of *Pp*XG5 products of XyG digestion with PACE. The assignment is enabled by XyG-related mutants. B, Structure of XyG and β-GGM in *mur3-3* or *mbgt1-1* mutants. C, Structure of XyG and β-GGM in *xxt1 xxt2* and *csla2* mutants. Bands in red dashed lines are XyG oligosaccharide dimers. M: Man; G: Glc. Markers M to $M_6$ are shown.

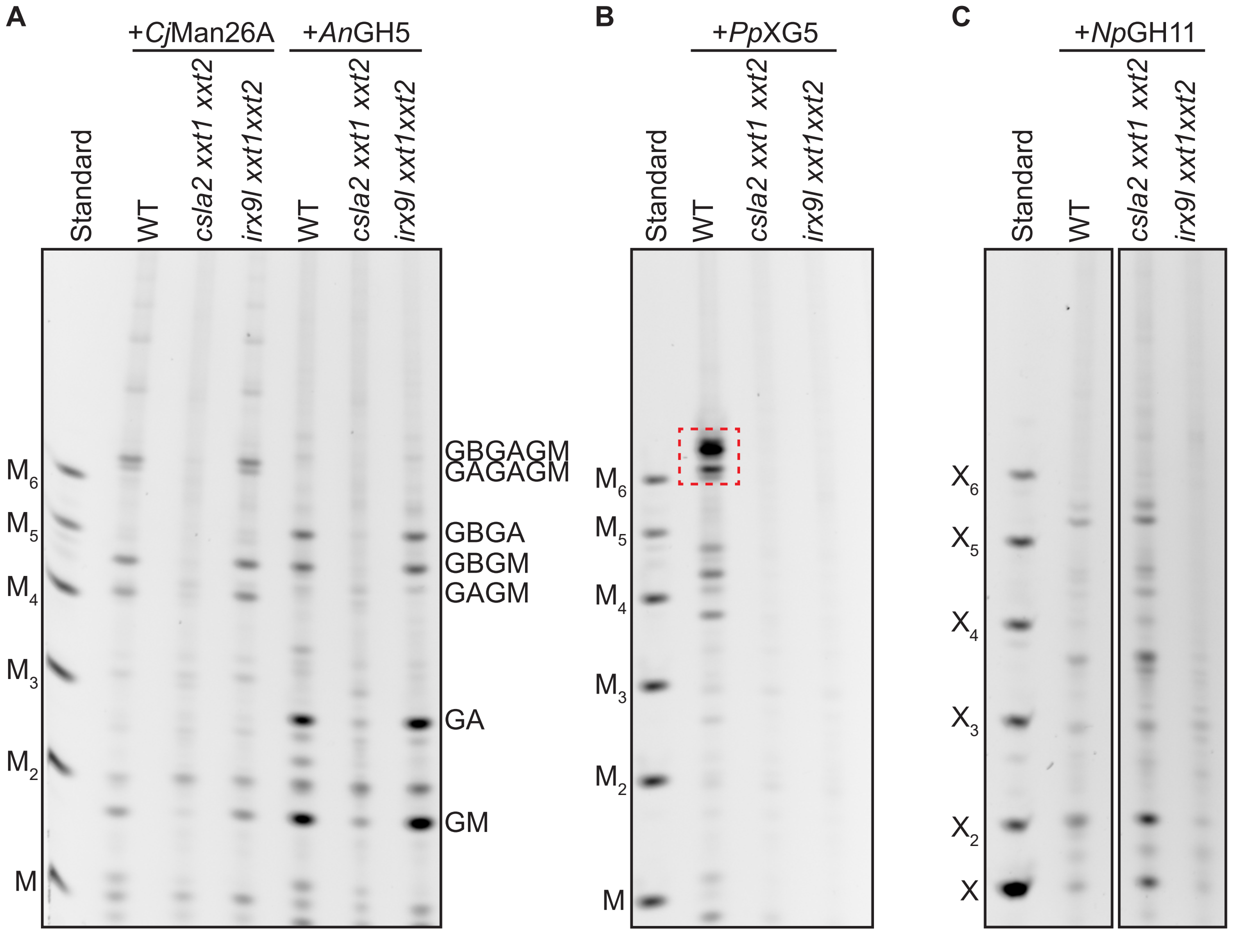

**Supplemental Figure S6. Arabidopsis callus hemicelluloses analyzed by PACE.** A, Callus mannan was analysed by both *Cj*Man26A and *An*GH5 mannanases. *csla2*-related mutants lack mannan accessed by *Cj*Man26A and *An*GH5. B, Callus XyG was analysed by *Pp*XG5. Bands in red dashed lines are XyG oligosaccharide dimers. C, Callus xylan was analyzed by *Np*GH11. *irx9l*-related mutants lack xylan accessed by *Np*GH11. M: Man; X: Xyl. Markers M to $M_6$ and X to $X_6$ are shown.

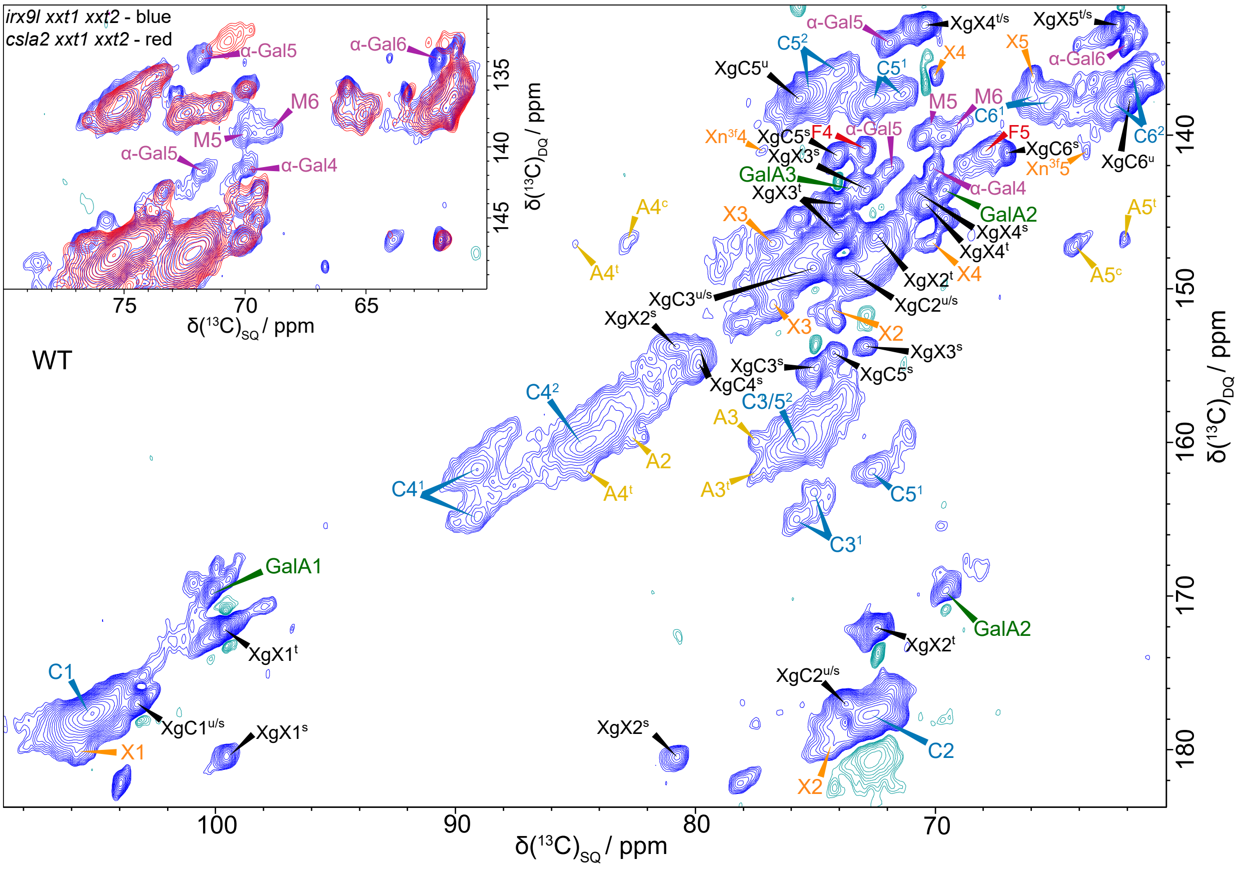

**Supplemental Figure S7. Solid-state NMR of WT Arabidopsis callus.** The carbohydrate region of a refocussed CP-INADEQUATE $^{13}$C MAS NMR spectrum of $^{13}$C enriched WT callus with the β-GGM Man peaks M5 and M6 labelled. Xyloglucan is also labelled as well as carbons in the major polysaccharides: galacturonic acid (GalA), terminal Xyl (X) and α-Gal and two arabinoses ($A^t$ and $A^c$). The terminal arabinose is labelled t and the other arabinose c. For cellulose, the environments have been split into two groups, domain 1 and 2 cellulose ($C^1$ and $C^2$). For xyloglucan, 5 sets of environments are seen depending on the substitution, labelled as unsubstituted backbone Glc ($XyC^u$), substituted backbone Glc ($XyC^s$), terminal Xyl on XyG ($XgX^t$), substituted Xyl ($XgX^s$), and Fuc (F). Assignments are listed in Supplementary Table 1. The inset shows an overlay for the M5, M6 region of a CP INADEQUATE spectrum of $^{13}$C enriched *irx9l xxt1 xxt2* callus (blue) with that of *csla2 xxt1 xxt2* callus. As expected, the β-GGM (M and α-Gal) peaks are missing from the *csla2 xxt1 xxt2* spectrum. Spectra were acquired at a $^{13}$C Larmor frequency of 213.8 MHz for WT callus and *csla2 xxt1 xxt2* and 251.6 MHz for *irx9l xxt1 xxt2*. The MAS frequency was 12.5 kHz and the spin-echo duration was 2.24 ms.

**Supplemental Table 1. Chemical shifts (in ppm) for different moieties obtained with solution or solid state NMR**

**Solution NMR------ $^{1}H$ and $^{13}C$ NMR chemical shifts for the GGM oligosaccharide**

| Name | Type, moiety | H1/C1 | H2/C2 | H3/C3 | H4/C4 | H5/C5 | H6/C6 | References |
|---|---|---|---|---|---|---|---|---|
| β-GGM | Substituted α-Gal | 5.2/99.3 | 3.94/78.5 | 4.16/74.3 | 3.76/70.8 | 3.5 | n/a | This work |
| β-GGM | β-Gal | 4.57/105.5 | 3.62/71.4 | 3.65/73.3 | 3.73/71.4 | n/a | n/a | This work |

**Solid State NMR------ $^{13}C$ NMR chemical shift of polysaccharides in WT callus**

| Name | Type, moiety | C1 | C2 | C3 | C4 | C5 | C6 | References |
|---|---|---|---|---|---|---|---|---|
| Spruce AcGGM | Man | 101.9 | 72.0 | n/a | 80.4 | 75.8 | 61.6 | Terrett et al., 2019 |
| Spruce AcGGM | Acetylated Man | 100.9 | 71.9 | 75.9 | 80.4 | 75.8 | 61.6 | Terrett et al., 2019 |
| Kiwi fruit GGM | Substituted Man | 100.4 | 70.9 | 70.4 | 78.1 | 69.6 | 68.6 | Schroder et al., 2001 |
| β-GGM | Man | 100.4 | 72.4 | 71.0 | 78.6 | 70.1 | 68.9 | This work |
| Kiwi fruit GGM | Un-substituted α-Gal | 99.2 | 71.9 | 71.5 | 76.9 | 69.6 | 60.4 | Schroder et al., 2001 |
| Kiwi fruit GGM | Substituted α-Gal | 98.9 | 78.1 | 70.4 | 69.6 | 75.5 | 61.3 | Schroder et al., 2001 |
| β-GGM | α-Gal | n/a | n/a | n/a | 70.1 | 71.9 | 62.0 | This work |
| Xyloglucan | Terminal α-Xyl | 99.5 | 72.4 | 74.0 | 70.2 | 62.0 | - | York et al., 1993 |
| Xyloglucan | Terminal α-Xyl | 99.7 | 72.5 | 74.4 | 70.5 | 62.3 | - | Dick-Pérez et al., 2011 |
| Xyloglucan | Terminal α-Xyl | 99.9 | 80.9 | 74.7 | 70.2 | 62.0 | - | York et al., 1993 |
| Xyloglucan | Terminal α-Xyl ($XgX^{t}$) | 99.6 | 72.5 | 74.0 | 70.4 | 62.4 | - | This work |
| Xyloglucan | Substituted α-Xyl ($XgX^{S}$) | 99.5 | 80.8 | 72.9 | 70.6 | 62.3 | - | This work |
| Xyloglucan | Un-substituted Glc | 105.0 | 72.7 | 75.9 | 83.0 | 73.8 | 61.5 | Dick-Pérez et al., 2011 |
| Xyloglucan | Substituted Glc | 103.1 | 73.7 | 75.0 | 80.5 | 74.4 | 67.7 | York et al., 1993 |
| Xyloglucan | Un-substituted Glc ($XgC^{u}$) | 103.3 | 73.7 | 75.0 | n/a | 75.7 | 61.9 | This work |
| Xyloglucan | Substituted Glc ($XgC^{S}$) | 103.3 | 73.7 | 75.0 | 79.8 | 74.0 | 67.0 | This work |
| Xyloglucan | L-Fucose | 100.7 | n/a | n/a | n/a | n/a | 17.2 | Watt et al., 1999 |
| Xyloglucan | L-Fucose | n/a | n/a | n/a | 72.8 | 67.8 | 16.8 | This work |
| Unknown | Terminal β-xylose | 105.8 | 74.4 | 76.8 | 70.0 | 65.9 | - | This work |
| Pectin | Galacturonic acid | 99.3 | 69.0 | 72.9 | n/a | n/a | n/a | This work |
| Arabinan | Terminal arabinose | 108.4 /110.0 | 82.2 | 77.7 | 84.9 | 62.3 | - | Wang et al., 2014 |
| Arabinan | Internal arabinose | 108.4 /110.0 | 82.2 | 77.8 | 82.5-83.2 | 64.0-67.9 | - | Wang et al., 2014 |
| Arabinan | Terminal arabinose ($A^{t}$) | 108.1 /107.6 | 81.9 | 77.4 | 84.8 | 61.9 | - | This work |
| Arabinan | Internal arabinose ($A^{c}$) | 109.9 | 82.1 | 77.4 | 82.6 | 63.9 | - | This work |

Figures in black are literature assignments shown for comparison. Figures in blue are assignments from the current work. n/a: not assigned.

**Supplemental Table 1. Chemical shifts (in ppm) for different moieties obtained with solution or solid state NMR (continued)**

| Solid State NMR------ $^{13}$C NMR chemical shift of polysaccharides in WT callus | | | | | | | | |
|---|---|---|---|---|---|---|---|---|
| Name | Type, moiety | C1 | C2 | C3 | C4 | C5 | C6 | References |
| Cellulose | Domain 1 (interior) | 105.2 | 72.4 | 74.5 | 88.9 | 72.7 | 65.0 | Dupree et al., 2015 |
| | Domain 2 (surface) | 105.2 | 72.4 | 74.5 | 84.2 | 75.5 | 62.7 | |
| Cellulose | Interior (type a-e) | 104.1-105.8 | 71.2-72.6 | 74.4-75.4 | 87.0-89.9 | 71.2-72.6 | 64.7-65.9 | Wang et al., 2016 |
| | Surface (type f, g) | 104.9-105.1 | 72.5-72.9 | 74.1-75.3 | 83.5-84.5 | 75.3 | 61.5-62.5 | |
| Cellulose | Domain 1 ($C^1$) | 105.3 | 72.5 | 75.7 | 89.1 | 72.6 | 65.1 | This work |
| | | 105.8 | 72.3 | 75.3 | 88.0 | 71.1 | 66.0 | |
| | | 104.9 | 72.3 | 75.3 | 89.2 | 71.4 | 65.7 | |
| | Domain 2 ($C^2$) | 104.8 | 72.6 | 75.5 | 83.2 | 73.8 | 61.4 | This work |
| | | 105.2 | 72.6 | 75.5 | 83.6 | 75.2 | 61.4 | |
| | | 105.8 | 72.6 | 75.5 | 84.4 | 75.7 | 61.7 | |

Figures in black are literature assignments shown for comparison. Figures in blue are assignments from the current work. n/a: not assigned.

| Supplemental Table S2. Plant species used for the phylogenetic tree | |
|---|---|
| **Plant Clade** | **Plant species** |
| Charophytes | *Klebsormidium nitens* |
| Bryophytes | *Marchantia polymorpha, Physcomitrella patens, Sphagnum fallax* |
| Lycophytes | *Selaginella moellendorffii* |
| Polypodiopsa (ferns) | *Azolla filiculoides, Salvinia cucullata* |
| Gymnosperms: | *Cycas micholitzii, Ginkgo biloba, Gnetum montanum, Picea abies, Picea glauca, Picea sitchensis, Pinus pinaster, Pinus sylvestris, Pinus taeda, Pseudotsuga menziesii, Taxus baccata* |
| Basal angiosperms | *Amborella trichopoda, Nymphaea colorata, Liriodendron chinense* |
| Non-Poaceae monocots | *Ananas comosus, Apostasia shenzhenica, Asparagus officinalis, Calamus simplicifolia, Elaeis guineensis, Musa acuminate, Phalaenopsis equestris, Spirodela polyrhiza, Zostera marina* |
| Poaceae | *Brachypodium distachyon, Cenchrus americanus, Hordeum vulgare, Lolium perenne, Miscanthus sinensis, Oropetium thomaeum, Oryza brachyantha, Oryza sativa ssp. Indica, Oryza sativa ssp. Japonica, Phyllostachys edulis, Saccharum spontaneum, Setaria italica, Sorghum bicolor, Triticum aestivum, Triticum turgidum, Zea mays B73, Zea mays B104, Zea mays PH207, Zoysia japonica ssp. Nagirizaki* |
| Dicots | *Actinidia chinensis, Amaranthus hypochondriacus, Aquilegia caerulea, Arabidopsis lyrata, Arabidopsis thaliana, Arachis ipaensis, Beta vulgaris, Brassica oleracea, Brassica rapa, Cajanus cajan, Capsella rubella, Capsicum annuum, Carica papaya,*<br>*Chenopodium quinoa, Cicer arietinum, Citrullus lanatus, Citrus clementina, Coffea canephora, Corchorus olitorius, Cucumis melo, Cucumis sativus L., Daucus carota,*<br>*Erythranthe guttata, Eucalyptus grandis, Fragaria vesca, Glycine max, Gossypium raimondii, Hevea brasiliensis, Malus domestica, Manihot esculenta, Medicago truncatula, Nelumbo nucifera, Petunia axillaris, Populus trichocarpa, Prunus persica, Pyrus bretschneideri, Ricinus communis, Schrenkiella parvula, Solanum lycopersicum, Solanum tuberosum, Tarenaya hassleriana, Theobroma cacao, Trifolium pratense, Utricularia gibba, Vigna radiata var. radiata,*<br>*Vitis vinifera, Ziziphus jujube* |

**Supplemental Table S4. Primers used in this study**

| Primer name | Sequence |
|---|---|
| Primers for identification of T-DNA insertion mutant | |
| SALK_065561-LP | CCCATCACTACCATCTTTCCC |
| SALK_065561-RP | TCATGCTCGAGGTGATATTCC |
| SAIL_852_F05C-LP | CCCATCACTACCATCTTTCCC |
| SAIL_852_F05C-RP | TCATGCTCGAGGTGATATTCC |
| SALK_065083-LP | AAAAAAGCATGTGTTGATGAACCC |
| SALK_065083-RP | GGTGGAAGTGGAGTGTCAAAG |
| SALK_071916-LP | CATTTTCACTAGATCCGCCAC |
| SALK_071916-RP | TCCGGTACAAGAACTGTAGCG |
| SALK_061576-LP | CAAGAACCAGACCGGTTGTATG |
| SALK_061576-RP | AGGTTGAAGAGGGGAAGTCTTG |
| SAIL_785_E02-LP | TAAACGTGTGTCCCCTAAACG |
| SAIL_785_E02-RP | AGAGAAATCTCGAGACCGGAC |
| SALK_101308-LP | TAAATTGTTTCCGCGGTACAC |
| SALK_101308-RP | AGTCACCAAAAGAACACGTGG |
| SALK_141953-LP | TGCAAACGAAATTAAACATAGGC |
| SALK_141953-RP | GAAGAAGAAACTGATTGGGGC |
| SALK_037323-LP | AAATAGGACGGTGGAGTGAG |
| SALK_037323-RP | ACTCTTGCATTCAGGGGAGG |
| MUR3-1-LP | CCAACCTACTTCCACCCAGC |
| MUR3-1 RP | GTTCTGATCTCGGGTCTGCG |
| GABI_552C10-LP | GGAAACCGCTCTTCCTACATC |
| GABI_552C10-RP | TCCGGTTTATCCGGTAAAAAC |
| Primers for amplification of MBGT1-3×Myc CDS | |
| MBGT1 forward | ATGCGACCCAAGAATTATTCTCAG |
| MBGT1 reverse | TCACAGATCTTCCTCAGAGA |

### Supplemental references:

1. **Dick-Pérez, M., Zhang, Y., Hayes, J., Salazar, A., Zabotina, O.A., and Hong, M.** (2011). Structure and interactions of plant cell-wall polysaccharides by two-and three-dimensional magic-angle-spinning solid-state NMR. Biochemistry-Us **50,** 989-1000.
2. **Schroder, R., Nicolas, P., Vincent, S.J., Fischer, M., Reymond, S., and Redgwell, R.J.** (2001). Purification and characterisation of a galactoglucomannan from kiwifruit (Actinidia deliciosa). Carbohydr Res **331,** 291-306.
3. **Terrett, O.M., Lyczakowski, J.J., Yu, L., Iuga, D., Franks, W.T., Brown, S.P., Dupree, R., and Dupree, P.** (2019). Molecular architecture of softwood revealed by solid-state NMR. Nat Commun **10,** 4978.
4. **Watt, D., Brasch, D., Larsen, D., and Melton, L.** (1999). Isolation, characterisation, and NMR study of xyloglucan from enzymatically depectinised and non-depectinised apple pomace. Carbohyd Polym **39,** 165-180.
5. **York, W.S., Harvey, L.K., Guillen, R., Albersheim, P., and Darvill, A.G.** (1993). The structure of plant-cell walls .36. Structural-analysis of tamarind seed xyloglucan oligosaccharides using beta-galactosidase digestion and spectroscopic methods. Carbohyd Res **248,** 285-301.
6. **Wang, T., Salazar, A., Zabotina, O.A., Hong, M.** (2014). Structure and dynamics of Brachypodium primary cell wall polysaccharides from two-dimensional $^{13}$C solid-state nuclear magnetic resonance spectroscopy. Biochemistry **53**, 2840-2854.
7. **Dupree, R., Simmons, T.J., Mortimer, J.C., Patel, D., Iuga, D., Brown, S.P. Dupree, P.** (2015). Probing the molecular architecture of Arabidopsis thaliana secondary cell walls using two- and three-dimensional $^{13}$C solid state nuclear magnetic resonance spectroscopy. Biochemistry **54**, 2335-2345.
8. **Wang, T., Yang, H., Kubicki, J.D., Hong, M.** (2016). Cellulose structural polymorphism in plant primary cell walls investigated by high-field 2D solid-state NMR spectroscopy and density functional theory calculations. Biomacromolecules **17**, 2210-2222.
